# Lung type II alveolar epithelial cells collaborate with CCR2^+^ inflammatory monocytes in host defense against an acute vaccinia infection in the lungs

**DOI:** 10.1101/2020.01.20.910927

**Authors:** Ning Yang, Joseph M. Luna, Peihong Dai, Yi Wang, Charles M. Rice, Liang Deng

## Abstract

The pulmonary immune system consists of a network of tissue-resident cells as well as immune cells that are recruited to the lungs during infection and/or inflammation. How the two immune components cross-talk during an acute viral infection is not well understood. Intranasal infection of mice with vaccinia virus causes lethal pneumonia and systemic dissemination. Here we report that vaccinia host range protein C7 is a critical virulence factor. Vaccinia virus with deletion of C7 (VACVΔC7L) is non-pathogenic in wild-type C57BL/6J mice, but it gains virulence in mice lacking STAT2, or IFNAR1, or MDA5/STING. We provide evidence that lung type II alveolar epithelial cells (AECs) provide first-line of defense against VACVΔC7L infection by inducing IFN-β and IFN-stimulated genes via the activation of the MDA5 and STING-mediated nucleic acid-sensing pathways. This leads to recruitment of CCR2^+^ inflammatory monocytes into the lungs to fight against viral dissemination.

## INTRODUCTION

Poxviruses are large cytoplasmic DNA viruses that are important human and veterinary pathogens. Smallpox, a highly contagious infectious disease with a high mortality rate that had claimed hundreds of millions of lives throughout the history, is caused by a human specific poxvirus--variola virus--through inhalation of airborne droplets. Prior to Edward Jenner’s vaccination using skin scarification with cowpox, variolation by inhalation of dried smallpox scabs was practiced in China as early as in the 10^th^ century to induce immunity against smallpox. Later, vaccinia virus became the vaccine strain of choice against smallpox and was used successfully throughout the world, which lead to smallpox eradication in 1980.

Studies on intranasal infection with vaccinia virus using mice may shed light on how the lung immune system defends against poxvirus infection. Extensive studies have been conducted to evaluate the pathogenicity of the virus and to define its virulence factors. By using genetic knockout mice or cell type depletion antibodies, several immune cell types and factors have been defined to play important roles in host defense against vaccinia infection. For example, antiviral CD8^+^ T cells and IFN-γ production by these cells are important for viral clearance (Goulding et al., 2014; Goulding et al., 2012). Batf3^−/−^ mice are more susceptible to VACV infection with more weight loss after infection, supporting a role of CD103^+^/CD8α^+^ DCs in cross-priming CD8^+^ T cells (Desai et al., 2018). In addition, cGAS^−/−^ mice are more susceptible to VACV infection, which suggests that cGAS-dependent DNA sensing is important for host defense against a DNA virus (Schoggins et al., 2014). All of these studies were performed using a low dose of wild type (WT) VACV infection because of the virulence nature of this virus.

Lung alveolar epithelial cells (AECs) provide both physical and biochemical barriers against respiratory infectious agents and provide first-line defense in an intranasal infection model. It has been generally accepted that lung type II AECs (AECII) respond to respiratory RNA virus infection (Stegemann-Koniszewski et al., 2016). However, the immunological response of lung AECII to a DNA virus infection has not been demonstrated.

In this study, we first demonstrated that vaccinia host range protein C7 is a virulence factor. VACVΔC7L is non-pathogenic in WT C57BL/6J mice at a high dose of infection intranasally, but gained virulence in STAT2, IFNAR1, or MDA5/STING-deficient mice. VACVΔC7L is non-pathogenic in RAG1-deficient mice, which lack T and B cells. Thus, this attenuated VACV mutant provides us with a model to evaluate the lung innate immune responses to acute pulmonary infection with a DNA virus. We found that VACVΔC7L infection triggers the release of IFN-β, CCL2, CXCL9, and CXCL10 into bronchioalveolar (BAL), whereas WT VACV infection fails to do so. Infection of primary lung AECII with VACVΔC7L *in vitro* leads to the induction of IFNB and IFN-stimulated genes (ISGs), which is dependent on the MDA5-dependent cytosolic dsRNA-sensing pathway as well as the STING-dependent cytosolic DNA-sensing pathway.

Given that both IFNAR1 and STAT2-deficient mice are more susceptible to VACVΔC7L infection, we hypothesize that type I IFN signaling plays an important role in restricting VACVΔC7L, either through stimulating IFNAR on the lung AEC and/or hematopoietic cell populations. To probe the relative contributions of lung non-hematopoietic resident cells versus hematopoietic cells in host defense against VACVΔC7L, we generated bone marrow chimeric mice. Our results indicate that the IFNAR signaling on the non-hematopoietic cells (likely the AECII) plays a critical role in host defense, and IFNAR signaling on hematopoietic cells also contributes. Using IFNAR1^fl/fl^-Sftpc^cre-ERT2^ mice, we showed that type I IFN signaling on lung AECIIs is important for host defense against VACVΔC7L infection.

To probe whether myeloid cells play a role in host defense against VACVΔC7L infection, we transiently depleted CCR2^+^ inflammatory monocytes in CCR2-DTR mice through administration of diphtheria toxin (DT), and found that depletion of CCR2^+^ inflammatory monocytes renders the mice susceptible to VACVΔC7L infection. Using CCR2-GFP mice, we found that after VACVΔC7L infection, CCR2^+^Ly6C^hi^ inflammatory monocytes are recruited into the lung parenchyma and differentiated into several populations, including Lyve1^−^ interstitial macrophages (IMs), Lyve1^+^ IMs, and DCs. Unlike WT mice, infection of MDA5/STING-deficient mice with VACVΔC7L fails to recruit Ly6C^+^ monocytes or to generate Lyve1^−^ IMs.

Taken together, we have developed a novel pulmonary infection model using attenuated vaccinia virus VACVΔC7L, which triggers MDA5/STING-dependent innate immunity in lung AECII cells. IFN-β production and its signaling on the AECII plays a critical role for host control of viral replication and dissemination. Our study also revealed the important role of CCR2^+^ inflammatory monocytes in host restriction of viral dissemination.

## RESULTS

### A mutant vaccinia virus lacking the C7L gene (VACVΔC7L) is highly attenuated in a murine intranasal infection model

To test whether the vaccinia host range protein C7 is a virulence factor, we generated a mutant vaccinia (Western Reserve) strain lacking the C7L gene through homologous recombination. The recombinant virus VACVΔC7L, which expresses mCherry under the vaccinia synthetic early/late promoter, is replication-competent in BSC40 cells (data not shown). We performed intranasal infection of WT VACV or VACVΔC7L in 6-8 week old WT C57BL/6J mice. WT VACV infection at 2 × 10^6^ pfu per mouse caused rapid weight loss and 100% lethality (**Figures 1A and 1B**). WT VACV infection at 2 × 10^5^ pfu per mouse resulted in an average of 22% weight loss, and 40% mortality (**Figures 1A and 1B**). By contrast, VACVΔC7L infection at the highest dose (2 × 10^7^ PFU) results in less than 20% weight loss, and all of the mice recovered their weight at 11 to 12 days post infection (**Figures 1C and 1D**). These results demonstrate that VACVΔC7L is attenuated by at least 100-fold compared with WT VACV in an intranasal infection model, and therefore C7 is a virulence factor.

**Figure 1.**
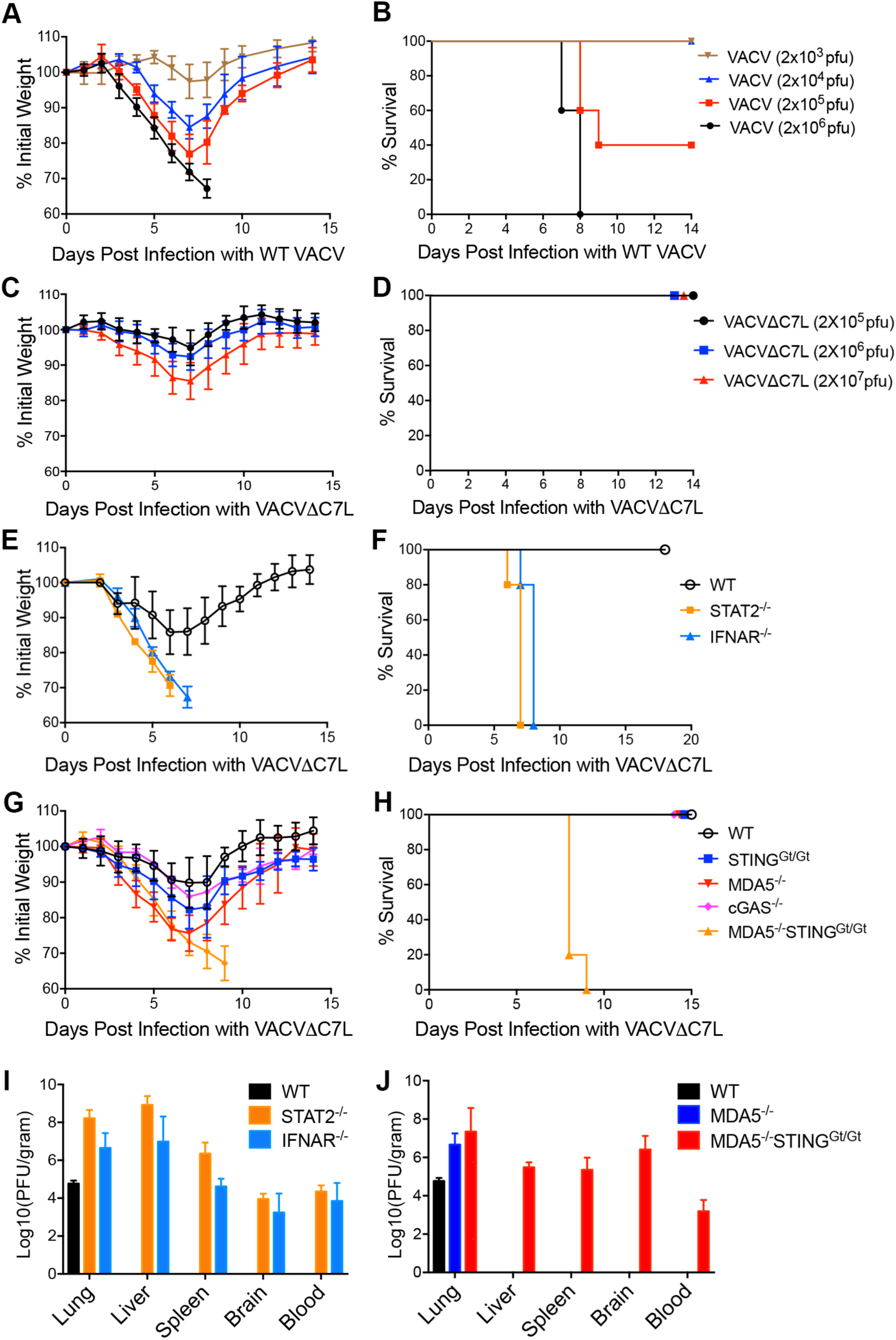
VACVΔC7L is highly attenuated in a murine intranasal infection model, but gains virulence in STAT2, IFNAR1-deficient, or MDA5^−/−^STING^Gt/Gt^ mice. (A) shown are the percentages of initial weight over days post intranasal infection with WT VACV at increasing doses. (B) Kaplan-Meier survival curve of WT C57BL/6J control mice (n=5 in each group) infected with WT VACV at increasing doses. (C) shown are percentages of initial weight over days post intranasal infection with VACVΔC7L at increasing doses. (D) Kaplan-Meier survival curve of WT C57BL/6J control mice infected with VACVΔC7L at increasing doses. (E) shown are the percentages of initial weight over days post intranasal infection with VACVΔC7L at a dose of 2 × 10^7^ pfu in STAT2^−/−^, IFNAR1^−/−^, or age-matched WT C57BL/6J control mice (n=5 in each group). A representative experiment is shown, repeated once. (F) Kaplan-Meier survival curve of STAT2^−/−^, IFNAR1^−/−^, or age-matched WT C57BL/6J mice infected with VACVΔC7L (n=5 in each group). (G) shown are the percentages of initial weight over days post intranasal infection with VACVΔC7L at a dose of 2 × 10^7^ pfu in cGAS^−/−^, STING^Gt/Gt^, MDA5^−/−^, MDA5^−/−^STING^Gt/Gt^ or age-matched WT C57BL/6J control mice (n=5 in each group). A representative experiment is shown, repeated once. (H) Kaplan-Meier survival curve of cGAS^−/−^, STING^Gt/Gt^, MDA5^−/−^, MDA5^−/−^STING^Gt/Gt^ or age-matched WT C57BL/6J control mice infected with VACVΔC7L (n=5 in each group). (I) Titers of VACVΔC7L in the lungs, livers, spleens, blood, and brains of STAT2^−/−^, IFNAR1^−/−^, or age-matched WT C57BL/6J control mice at day 4 post intranasal infection with VACVΔC7L at a dose of 2 × 10^7^ pfu. Data are represented as mean ± SEM (n=3-5). (J) Titers of VACVΔC7L in the lungs, livers, spleens, blood, and brains of MDA5^−/−^, MDA5^−/−^ STING^Gt/Gt^ or age-matched WT C57BL/6J control mice at day 4 post intranasal infection with VACVΔC7L at a dose of 2 × 10^7^ pfu. Data are represented as mean ± SEM (n=3-5).

### Type I IFN signaling is crucial for host control of VACVΔC7L infection in the lungs

To probe the mechanism of attenuation, we first tested whether type I IFN signaling is important for host defense against VACVΔC7L. We intranasally infected WT, STAT2^−/−^, or IFNAR1^−/−^ mice with VACVΔC7L at a dose of 2 × 10^7^ pfu and monitored weight loss and survival of the mice over time. We found that in contrast to WT mice, the STAT2^−/−^ and IFNAR^−/−^ mice were highly susceptible to VACVΔC7L infection, with rapid weight loss, severe illness, and death (**Figures 1E and 1F**). The median survival times for STAT2^−/−^ and IFNAR1^−/−^ mice were 7 days and 8 days, respectively (**Figure 1F**). We compared the viral titers in various tissues from WT, STAT2^−/−^, or IFNAR1^−/−^ mice at day 4 post infection with VACVΔC7L at 2 × 10^7^ pfu. We found that VACVΔC7L infection of WT mice caused localized infection in the lungs without dissemination of the virus or viremia (**Figure 1I**). VACVΔC7L infection caused higher viral titers in the lungs of STAT2^−/−^ or IFNAR1^−/−^ mice compared with those in the WT mice (**Figure 1I**). We also observed viremia and dissemination of the virus to various distant organs including livers, spleens, and brains in STAT2^−/−^ and IFNAR1^−/−^ mice (**Figure 1I**).

To determine the LD50 of VACVΔC7L virus in STAT2^−/−^ or IFNAR1^−/−^ mice, we intranasally infected these mice with decreasing doses of VACVΔC7L. Our results show that the LD50 of VACVΔC7L in STAT2^−/−^ and IFNAR1^−/−^ mice is around 1000 pfu (**Figures S1A-S1D**).

### Both the cytosolic dsRNA and DNA-sensing pathways plays important roles in restricting pulmonary VACVΔC7L infection

We have previously shown that the cytosolic DNA- and dsRNA-sensing pathways are important for host immune detection of vaccinia infection, leading to type I IFN production in dendritic cells (DCs) and epithelial cells in a cell type-specific manner (Dai et al., 2014; Deng et al., 2008). cGAS^−/−^ mice are highly suspectible to WT VACV infection (Schoggins et al., 2014). We performed intranasal infection of VACVΔC7L at a dose of 2 × 10^7^ pfu in cGAS^−/−^, STING^Gt/Gt^, MDA5^−/−^, or MDA5^−/−^STING^Gt/Gt^ mice and found that the average percentages of weight loss were 10%, 14%, 18%, 24%, and 27% for WT, cGAS^−/−^, STING^Gt/Gt^, MDA5^−/−^ and MDA5^−/−^STING^Gt/Gt^ mice, respectively, at day 7 post infection. All of the WT, cGAS^−/−^, STING^Gt/Gt^, or MDA5^−/−^ mice subsequently gained weight and recovered from acute illness. By contrast, all of the MDA5^−/−^STING^Gt/Gt^ mice died at day 8 or 9 post infection (**Figures 1G and 1H**). We compared viral titers in various organs harvested from WT, MDA5^−/−^ and MDA5^−/−^ STING^Gt/Gt^ mice and found that although viral titers were higher in the infected lungs of MDA5^−/−^ mice compared with those in WT mice, VACVΔC7L infection was confined to the lungs in MDA5^−/−^ mice. By contrast, in MDA5^−/−^STING^Gt/Gt^ mice infected with VACVΔC7L, we observed systemic dissemination of the virus at day 4 post infection (**Figure 1J**). These results are consistent with the differences in mortality in MDA5^−/−^ and MDA5^−/−^STING^Gt/Gt^ mice upon VACVΔC7L infection (**Figure 1H**). Based on these results, we conclude that both the cytosolic dsRNA-sensing pathway mediated by MDA5 and the DNA-sensing pathway mediated by cGAS/STING play important roles in host restriction of vaccinia infection in the lungs and in preventing systemic dissemination.

### Intranasal infection of VACVΔC7L leads to the production of type I IFN and proinflammatory cytokines and chemokines in the infected lungs

We hypothesize that intranasal infection with VACVΔC7L induces lung innate immunity including type I IFN production whereas WT vaccinia virus does not. To test that, we isolated bronchoalveolar lavage fluid (BAL) at day 1 and day 3 post infection with either WT VACV or VACVΔC7L at 2 × 10^7^ pfu. We found that at day 1 post infection, VACVΔC7L induced detectable IFN-β level in BAL, whereas WT VACV did not. At day 3 post infection, VACVΔC7L induced much higher levels IFN-β levels in the BAL compared with those collected at day 1 post infection, whereas WT VACV had a very small induction of IFN-β level at day 3 post infection (**Figures 2A and 2B**). Luminex analysis of proinflammatory cytokines and chemokines in the BAL showed that VACVΔC7L infection resulted in the release of IL-6, Ccl2, IFN-γ, CXCL10, CXCL9 into the BAL, whereas both WT VACV and VACVΔC7L induced VEGF release (**Figures 2A and 2B**).

**Figure 2.**
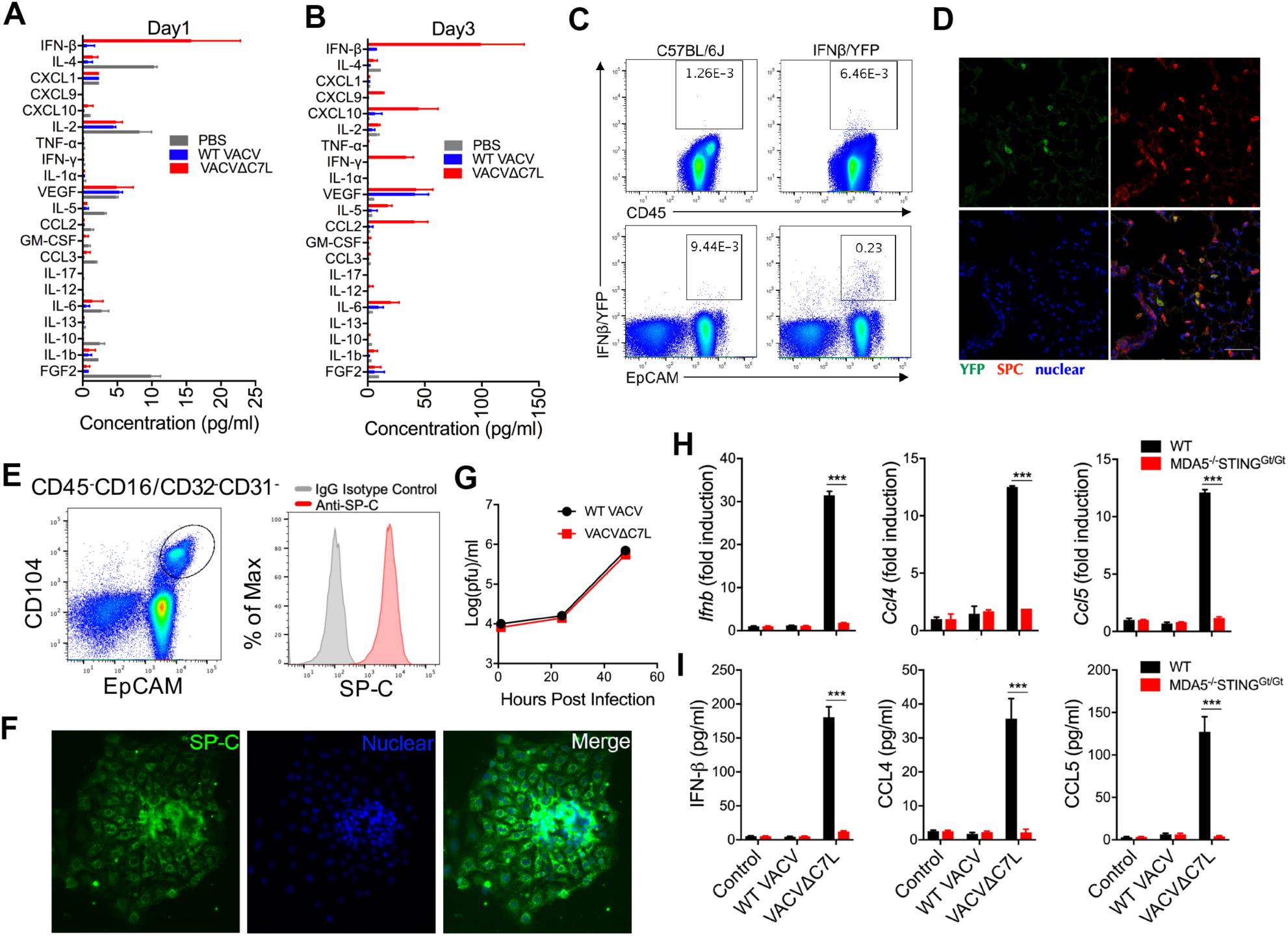
Lung AECIIs produce IFN-β and proinflammatory cytokines and chemokines upon VACVΔC7L infection in a MDA5/STING-dependent manner. (A) and (B) Levels of IFN-β and other cytokines and chemokiens in BAL from VACVΔC7L or WT VACV-infected mice collected at day1 and day 3 post infection determined by ELISA or Luminex. (C) Dot plots showing percentages of IFNβ/YFP positive cells among CD45^+^ immune cells and CD45^−^EpCAM^+^ lung AECIIs in VACVΔC7L-infected lungs from IFNβ-YFP and WT C57BL/6J mice determined by FACS. (D) Immunohistology of lung section from IFNβ-YFP mice infected with VACVΔC7L collected at 1 day post infection. Top left: IFNβ-YFP^+^ cells (green); Top right: surfactant protein C (SPC) positive AECII (red); Bottom left: DAPI staining of nuclei (blue); Bottom right: overlay of the three colors showing co-localization of green and red signals demonstrating that lung AECIIs are IFN-β producing cells. (E) Gating strategy for the isolation of lineage negative epithelial progenitor cells that are CD45^−^ CD16/CD32^−^CD31^−^EpCAM^+^CD104^+^. Cells were cultured in vitro on Matrigel-coated plates as described in methods for 4-5 days. The identify of AECII cells were confirmed by SPC^+^ staining determined by FACS. (F) Immunofluorescence staining of SPC of *in vitro* cultured AECII cells. (H) RT-PCR analyses of Ifnb, Ccl4, and Ccl5 gene expression of AECII cells from WT or MDA5^−/−^STING^Gt/Gt^ mice infected with either WT VACV or VACVΔC7L at a MOI of 10. (I) ELISA analyses of IFN-β, CCL4, CCL5 levels in the supernatants of AECII culture from WT or MDA5^−/−^STING^Gt/Gt^ mice infected with either WT VACV or VACVΔC7L at a MOI of 10.

### Lung type II alveolar epithelial cells (ACEIIs) are the major producers of IFN-β *in vivo* at an early phase of intranasal infection of VACVΔC7L

The lung epithelial cells and alveolar macrophages (AMs) provide the first line defense against pulmonary pathogen infection. To determine which cell population(s) are the major producer(s) of IFN-β upon VACVΔC7L infection, we used IFN-β-yellow fluorescent protein (YFP) knockin mice to map the cell type(s) responsible for IFN-β production induced by VACVΔC7L infection. We found that the majority of IFN-β/YFP positive cells are CD45^−^EpCAM^+^ (**Figure 2C**). Confocal microscopy of lung sections from IFN-β/YFP at day one post infection with VACVΔC7L showed IFN-β/YFP-positive cells that overlap with lung AECII marker *surfactant protein C* (*SP-C*) (**Figure 2D**). Although lung AMs can be infected with either WT VACV or VACVΔC7L in vivo (data not shown), infection of AMs with either WT VACV or VACVΔC7L *in vitro* does not result in IFNB gene induction or IFN-β production (**Figures S2A and S2B**). To test whether AMs can respond to poxvirus infection, we infected them with a highly attenuated modified vaccinia virus Ankara (MVA), which is non-replicative in most mammalian cells. We have previously shown that MVA infection in conventional dendritic cells (cDCs) induces type I IFN production via the cGAS/STING/IRF3-mediated cytosolic DNA-sensing pathway (Dai et al., 2014). We found that MVA infection of AMs induces IFNB gene expression and IFN-β production (**Figures S2A and S2B**). and MVAΔC7L induces higher levels of IFNB gene expression and protein secretion compared with MVA (**Figures S2A and S2B**). These results provide evidence to support that vaccinia C7 plays an inhibitory role of the IFN production pathway and also suggest that there are additional vaccinia inhibitors of the cGAS/STING/IRF3 pathway that prevent the induction of IFNB in AMs by VACVΔC7L.

### Primary murine lung AECIIs induces Ifnb, Ccl4, and Ccl5 gene expression and protein secretion in a MDA5/STING-dependent manner upon VACVΔC7L infection

To firmly establish that the lung AECIIs are capable of producing IFN-β and proinflammatory chemokines upon VACVΔC7L infection and to test whether the induction is dependent on the MDA5 and STING-mediated cytosolic dsRNA and DNA-sensing pathways, we isolated the lineage negative epithelial progenitor cells, CD45^−^CD16/CD32^−^CD31^−^EpCAM^+^CD104^+^, from the lungs of WT and MDA5^−/−^STING^Gt/Gt^ mice. The cells were cultured *in vitro* to allow differentiation into AECIIs. *SP-C* expression was determined by FACS analysis as well as immunofluorescence staining to confirm AECII identity (**Figures 2E and 2F**). We next tested the replication capacities of WT VACV and VACVΔC7L in primary lung AECII and found that they were similar (**Figure 2G**). To test the innate immune responses of lung AECIIs to WT VACV or VACVΔC7L infection, AECII from WT and MDA5^−/−^STING^Gt/Gt^ mice were infected with either WT VACV or VACVΔC7L at a multiplicity of infection (MOI) of 10. The cells were collected at 12 h post infection and quantitative PCR analyses were performed. We found that infection of WT lung epithelial cells with VACVΔC7L induced higher levels of expression of Ifnb, Ccl4, and Ccl5 compared with WT VACV (**Figure 2H**). ELISA analysis of supernatants collected at 24 h post infection with either WT VACV or VACVΔC7L showed that VACVΔC7L infection resulted in the secretion of IFN-β, CCL4 and CCL5 by WT AECIIs. By contrast, the MDA5^−/−^STING^Gt/Gt^ AECII failed to induce Ifnb, Ccl4, and Ccl5 gene expression and to produce IFN-β, CCL4 and CCL5 upon VACVΔC7L infection (**Figures 2H and 2I**). However, MDA5-deficient AECIIs had modest reduction of IFNB gene expression and IFN-β secretion compared with WT cells in response to VACVΔC7L and STING-deficient AECII had similar capacitites to induce IFNB in respsone to VACVΔC7L (**Figures S2C and S2D**). These results demonstrate that the induction of Ifnb, Ccl4, and Ccl5 gene expression and protein secretion by VACVΔC7L is dependent on both MDA5 and STING.

### Vaccinia C7 inhibits IFNB gene induction by innate immune pathways and type I IFN signaling

To understand the mechanism by which vaccinia C7 antagonizes the IFN pathway, we utilized a dual-luciferase assay system to evaluate the role of vaccinia C7 in the regulation of STING, TBK1, MAVS, TRIF, TLR3, or IRF3-induced IFNB promoter activation in HEK293T cells. HEK293T cells were transfected with plasmids expressing IFNB-firefly luciferase reporter, a control plasmid pRL-TK that expresses *Renilla* luciferase, innate immune sensor/adaptor, and vaccinia C7L, as indicated. Dual luciferase assays were performed at 24 h post transfection. The relative luciferase activity was expressed as arbitrary units by normalizing firefly luciferase activity to Renilla luciferase activity. Overexpression of STING resulted in a 30-fold induction of IFNB promoter activity compared with that in the control sample without STING. Co-transfection of increasing amounts of C7L expression plasmid led to a significant reduction of STING-induced IFNB promoter activity (**Figure 3A**). Similarly, overexpression of TBK1 resulted in a 400-fold induction of IFNB promoter activity compared with the control. Co-transfection of increasing amounts of C7L expression plasmid led to an over 90% reduction of TBK1-induced IFNB promoter activity (**Figure 3B**). The TBK1-IRF3 axis is important for signal transduction in several pathways, including cGAS-cGAMP-STING, RIG-I/MDA5-MAVS, TLR3-TRIF. MAVS or TRIF overexpression induced a 500-fold induction of IFNB promoter activity compared with the control. C7 blocked the MAVS or TRIF-induced luciferase signal by 70% (**Figure 3C and 3D**). Transfection of TLR3 and treatment with poly I:C resulted in a 9-fold induction of IFNB promoter activity compared with an empty vector control (**Figure 3E**). Overexpression of C7 resulted in the reduction of poly (I:C)/TLR3-induced IFNB promoter activity by up to 90% (**Figure 3E**). These results indicate that overexpression of C7 in HEK293T cells exerts an inhibitory effect on STING, MAVS, TLR3/poly (I:C), TRIF, and TBK1-induced IFNB promoter activity. IRF3 is a member of the interferon regulatory transcription factor (IRF) family and it is an essential transcription factor for the IFNB promoter. Since TBK1/IRF3 is a common node in these diverse DNA- and RNA-sensing pathways, it is possible that C7 targets the step that leads to the activation of IRF3, resulting in the failure of IRF3 phosphorylation. We found that over-expression of C7 caused a 70% reduction of IRF3-induced IFNB promoter activity (**Figure 3F**), whereas overexpression of C7 failed to reduce IRF3-5D-induced IFNB promoter activity (**Figure 3G**). IRF3-5D is a constitutive active, phosphorylation-mimetic mutation of IRF3. In addition, we found that C7 does not affect NFKB gene activation induced by TRIF overexpression (**Figure 3H**). Taken together, our results indicate that C7 functions through inhibition of IRF3 activation.

**Figure 3.**
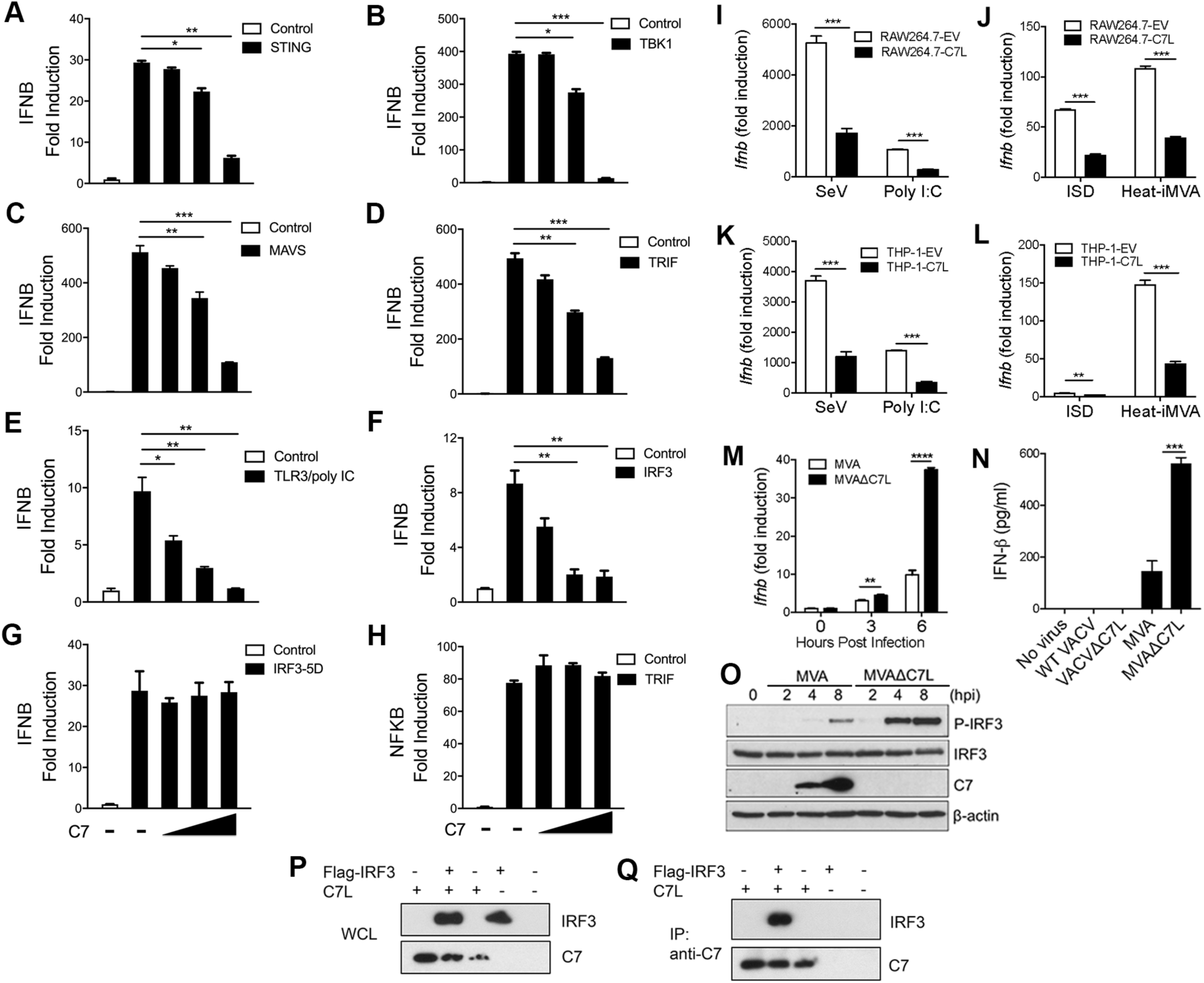
Vaccinia C7 inhibits IFNB gene induction by innate immune pathways by interacting with IRF3 and preventing IRF3 phosphorylation. (A) Dual-luciferase assay of HEK293T cells transfected with IFNB-firefly luciferase reporter, a control plasmid pRL-TK expressing *Renilla* luciferase, vaccinia C7L-expressing or control plasmid, and STING-expressing plasmid. Cells were harvested at 24 h post transfection. Data are represented as mean ± SEM. (B) Luciferase assay was carried out in the same condition as in (A) except that TBK1-expressing plasmid was used instead of STING-expressing plasmid. (C) Luciferase assay was carried out in the same condition as in (A) except that MAVS-expressing plasmid was used. (D) Luciferase assay was carried out in the same condition as in (A) except that TRIF-expressing plasmid was used. (E) Luciferase assay was carried out in the same condition as in (A) except that TLR3-expressing plasmid was used. 24 h post transfection, cells were treated with poly I:C for another 24 h before harvesting. (F) Luciferase assay was carried out in the same condition as in (A) except that IRF3-expressing plasmid was used. (G) Luciferase assay was carried out in the same condition as in (A) except that IRF3-5D-expressing plasmid was used. (H) Dual-luciferase assay of HEK293T cells transfected with NFkB-firefly luciferase reporter, a control plasmid pRL-TK expressing *Renilla* luciferase, vaccinia C7L-expressing or control plasmid, and TRIF-expressing plasmid. Cells were harvested at 24 h post transfection. Data are represented as mean ± SEM. (I) RAW264.7 stable cell line expressing vaccinia C7 (RAW264.7-C7L) or with empty vector (RAW264.7-EV) were infected with Sendai virus (SeV) or treated with poly I:C. Cells were collected 24 h later. IFNB gene expression level was measured by quantitative real-time PCR. Data are represented as mean ± SEM. (J) RAW264.7-C7L or RAW264.7-EV were transfected with interferon stimulatory DNA (ISD) or infected with heat-inactivated MVA (Heat-iMVA). Cells were collected 24 h later. IFNB gene expression level was measured. (K) THP-1 stable cell line expressing vaccinia C7 (THP-1-C7L) or with empty vector (THP-1-EV) were infected with Sendai virus (SeV) or treated with poly I:C. Cells were collected 24 h later. IFNB gene expression level was measured. Data are represented as mean ± SEM. (L) THP-1-C7L or THP-1-EV were transfected with ISD or infected with Heat-iMVA. Cells were collected 24 h later. IFNB gene expression level was measured. (M) Bone marrow-derived dendritic cells (BMDCs) were infected with either MVA or MVAΔC7L at a MOI of 10. Cells were collected at 3 h and 6 h post infection. The IFNB gene expression levels were determined by quantitative PCR analyses. Data are represented as mean ± SEM. (N) BMDCs were infected with either WT VACV, VACVΔC7L, MVA, or MVAΔC7L at a MOI of 10. Supernatants were collected at 22 h post infection. The IFN-β levels in the supernatants were determined by ELISA. (O) Western blot analyses of lysates from MVA or MVAΔC7L infected BMDCs. Cells were collected at 2, 4, and 8 h post infection. Anti-phospho-IRF3, -IRF3, -β-actin, and -C7 antibodies were used. A representative experiment is shown, repeated once. (P) HEK293T cells were co-transfected with Flag-tagged IRF3 or C7L either alone or in combination. The whole cell lysates were blotted with anti-Flag and anti-C7 antibody. (Q) Same as in (P). The whole cell lysates were immunoprecipitated with anti-C7 antibody, and immunoblotted with anti-Flag antibody.

### Overexpression of vaccinia C7 in immune cells inhibits IFNB gene induction and IFN-β signaling

To assess the effect of vaccinia C7 in IFNB gene induction in immune cells, we generated two cell lines stably expressing vaccinia C7, including murine macrophage RAW264.7 and human THP-1. An empty vector with a drug selection marker was also used to generate a control cell line. THP-1 stable cell line expressing C7 or with an empty vector were differentiated by phorbol-12-myristate-13-acetate (PMA) for 3 days before they were used for the experiments. The cells were either infected with Sendai virus (SeV), or heat-inactivated MVA (Heat-iMVA), or they were incubated with poly I:C, or transfected with interferon stimulatory DNA (ISD), a 45-bp non-CpG oligomer from *Listeria monocytogenes*. After 24 h, the IFNB gene expression level was measured by quantitative real-time PCR. SeV infection induced the highest level of IFNB gene expression in both RAW264.7 and THP-1 cells among all of the stimuli used in this experiment, and the overexpression of vaccinia C7 resulted in the reduction IFNB gene expression by 67% and 68%, respectively (**Figure 3I**, **3K**). Vaccinia C7 also attenuated poly (I:C)-induced IFNB gene expression in RAW264.7 and THP-1 cells by 73% and 75%, respectively (**Figure 3I**, **3K**). Similarly, vaccinia C7 reduced Heat-iMVA-induced IFNB gene expression in RAW264.7 and THP-1 cells by 64% and 71%, respectively (**Figure 3J**, **3L**). Furthermore, vaccinia C7 reduced ISD-induced IFNB gene expression in RAW264.7 by 68% (**Figure 3J**). SeV is a negative-sense, single-stranded RNA virus that belongs to the paramyxoviridae family. SeV infection can be sensed by the cytoplasmic RNA sensors, including retinoic-acid inducible gene-I (RIG-I) and melanoma differentiation-associated gene 5 (MDA-5) (Gitlin et al., 2010; Kawai et al., 2005). This can lead to the activation of the MAVS/TBK1/IRF3 axis. Poly (I:C) activates the endosomal dsRNA sensor, TLR3, which leads to activation of the TRIF/TBK1/IRF3 axis. Heat-iMVA activates the cytosolic DNA-sensor cGAS, which leads to the generation of the second messenger, cyclic GMP-AMP (cGAMP), and the activation of the STING/TBK1/IRF3 axis (Dai et al., 2017). Taken together, these results indicate that vaccinia C7 inhibits multiple innate immune sensing pathways in macrophages.

### Vaccinia C7 downregulates IRF3 phosphorylation in BMDCs induced by MVA

We found that similar to infection in lung alveolar macrophages, MVAΔC7L induced higher levels of IFNB gene expression compared with MVA in BMDCs (**Figure 3M**). Whereas neither WT VACV nor VACVΔC7L induced IFN-β secretion in BMDCs, both MVA and MVAΔC7L infection triggered IFN-β production, with MVAΔC7L inducing higher levels of IFN-β secretion in BMDCs compared with MVA (**Figure 3N**). Western blot analysis showed that MVAΔC7L infection of BMDCs also induced higher levels of phosphorylation of IRF3 compared with MVA, indicating that C7 might block IRF3 phosphorylation (**Figure 3O**).

### Vaccinia C7 interacts with IRF3

To probe the mechanisms by which vaccinia C7 exerts its inhibitory effects on IRF3 phosphorylation, a co-immunoprecipitation assay was performed to determine whether vaccinia C7 interacts with IRF3. HEK293T cells were co-transfected with Flag-tagged IRF3 or C7 either alone or in combination. The whole cell lysates (WCL) were prepared and blotted with anti-FLAG and anti-C7 antibodies, which showed the expression of IRF3 and C7 in transfected cells (**Figure 3P**). Following immunoprecipitation of the whole cell lysates with an anti-C7 antibody, the C7-interacting proteins were then probed with an anti-Flag antibody. We observed that the Flag-tagged IRF3 was pulled down by the anti-C7 antibody from whole cell lysates (**Figure 3Q**). Taken together, our results show that C7 interacts with IRF3 to mediate its inhibitory effects on IFN gene induction.

### Type I IFN signaling on lung non-hematopoietic resident cells plays a critical role in host defense against VACVΔC7L infection

VACVΔC7L gains virulence in IFNAR1^−/−^ or STAT2^−/−^ mice in an intranasal infection model, which indicates that type I IFN signaling is crucial for controlling VACVΔC7L infection. To distinguish the contributions of IFNAR signaling in hematopoietic cells vs. non-hematopoietic cells to host restriction of VACVΔC7L infection in the lungs, we generated bone marrow chimeras and infected them with VACVΔC7L intranasally at 2 × 10^7^ pfu. Analysis of CD45.1 and CD45.2 markers of immune cells in the bone marrow chimeras showed the desired reconstitution of hematopoietic cells in the blood (**Figure S3A-S3D**). VACVΔC7L infection in WT → WT mice resulted in transient weight loss and all of the mice survived the infection. By contrast, VACVΔC7L infection in Ifnar1^−/−^ → Ifnar1^−/−^ mice resulted in rapid weight loss and 100% mortality (**Figure 4A and 4B**). All of the Ifnar1^−/−^ recipient mice reconstituted with WT bone marrow cells succumbed to VACVΔC7L infection, indicating that type I IFN signaling on non-hematopoietic resident cells are important for host restriction of VACVΔC7L infection (**Figure 4A and 4B**). By contrast, all of the WT recipient mice reconstituted with Ifnar1^−/−^ bone marrow cells survived despite losing more weight compared with WT recipient mice reconstituted with WT bone marrow cells. This suggests that type I IFN signaling on hematopoietic resident cells contribute to host defense against VACVΔC7L but with a limited capacity (**Figure 4A and 4B**).

**Figure 4.**
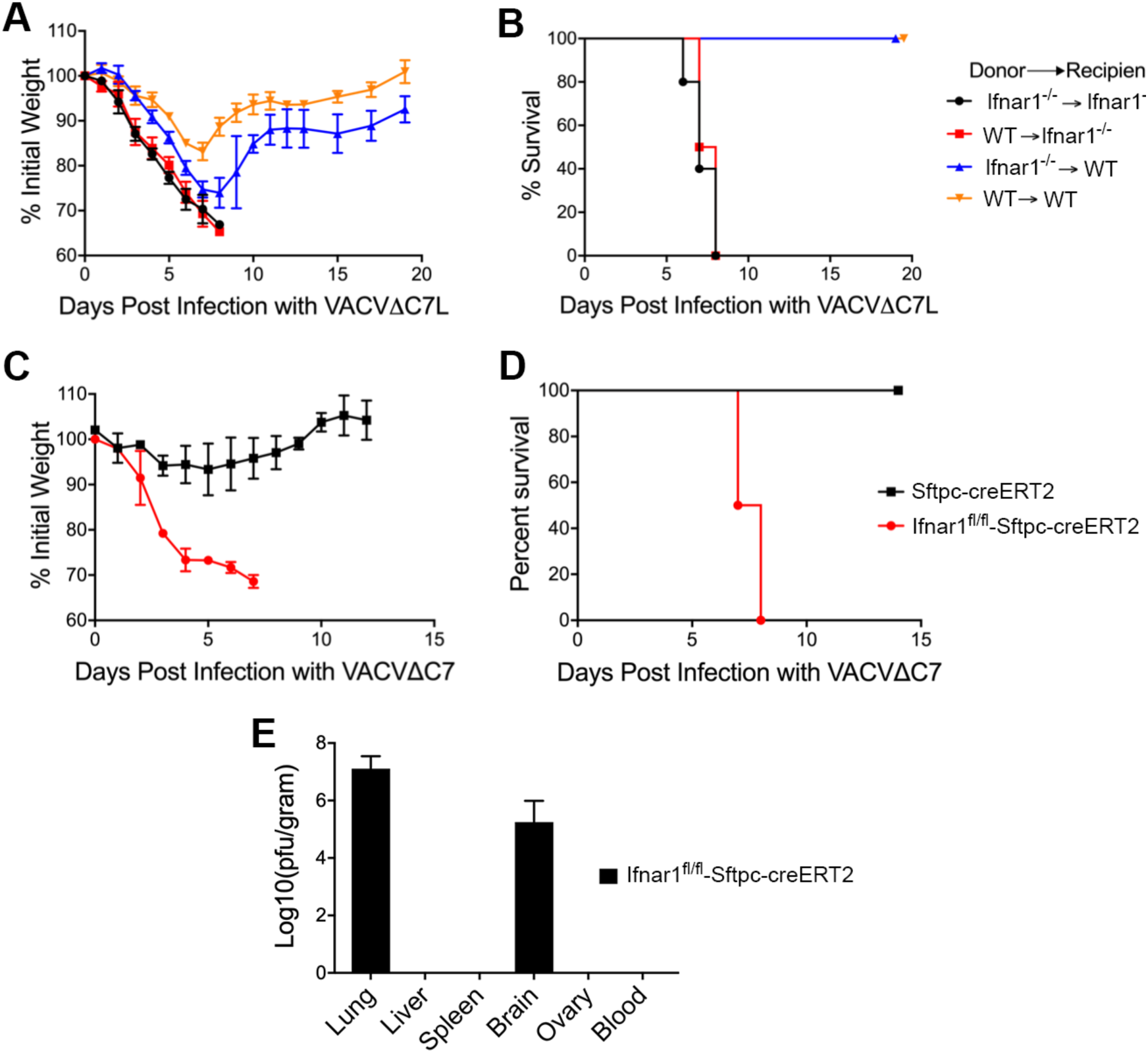
Type I IFN signaling in lung non-hemopoietic resident cells plays crucial role in host retriction of vaccinia infection. (A) shown are the percentages of initial weight over days post intranasal infection with VACVΔC7L at 2 × 10^7^ pfu in bone marrow chimeras (Ifnar1^−/−^→ Ifnar1^−/−^, WT → WT, Ifnar1^−/−^→ WT, and WT→ Ifnar1^−/−^). (B) Kaplan-Meier survival curve of bone marrow chimeras (Ifnar1^−/−^→ Ifnar1^−/−^, WT → WT, Ifnar1^−/−^→ WT, and WT→ Ifnar1^−/−^) infected with VACVΔC7L at 2 × 10^7^ pfu (n=5 in each group). A representative experiment is shown, repeated once. (C) shown are the percentages of initial weight over days post intranasal infection with VACVΔC7L at 2 × 10^7^ pfu in Ifnar1^fl/fl^-Sftpc^creERT2^ and Sftpc^creERT2^ mice treated with tamoxifen. (D) Kaplan-Meier survival curve of tamoxifen-treated Ifnar1^fl/fl^-Sftpc^creERT2^ and Sftpc^creERT2^ mice infected with VACVΔC7L at 2 × 10^7^ pfu (n=5 in each group). A representative experiment is shown, repeated once. (E) Titers of VACVΔC7L in the lungs, livers, spleens, blood, and brains of tamoxifen-treated Ifnar1^fl/fl^-Sftpc^creERT2^ at day 7 or 8 post intranasal infection with VACVΔC7L at a dose of 2 × 10^7^ pfu. Data are represented as mean ± SEM (n=3-5).

### Lung AECIIs are crucial targets of type I IFN induced by VACVΔC7L intranasal infection

Given that lung AECIIs are the major early producers of IFN-β upon intranasal VACVΔC7L infection and type I IFN signaling on non-hematopoietic resident cells plays a crucial role in controlling VACVΔC7L infection, we hypothesized that IFN signaling on lung AECII is important for restricting VACVΔC7L infection in the lungs. To test that, we used Ifnar1^fl/fl^-Sftpc^creERT2^ mice, which lacks IFNAR1 specifically in lung AECII upon tamoxifen-induced cre expression (Rock et al., 2011). Whereas the control mice had mild transient weight loss upon VACVΔC7L infection, all of the Ifnar1^fl/fl^-Sftpc^creERT2^ mice suffered severe weight loss and were euthanized at day 7 or 8 post infection when they lost more than 30% of their original weight (**Figure 4C and 4D**) Viral titers in various organs including blood were determined from these animals. We only detected high viral titers in the lungs and brains, but not in the liver, spleen, ovaries, or blood (**Figure 4E**). These results indicate that type I IFN signaling in lung AECII contributes to eradicating viral infection in the lungs as well as to controlling neurovirulence of vaccinia virus.

### Intranasal administration of IFN-β rescues mice from lethal WT VACV infection

We reasoned that if the inability to induce IFN-β from lung AECII by WT VACV is the main contributing factor for its virulence, we should be able to rescue mice from lethal infection with WT VACV. To test that, we infected 6-8 week old WT C57BL/6J mice with WT VACV at 2 × 10^5^ pfu or 2 × 10^6^ pfu. They were either treated with intranasal administarion of IFN-β (1 µg per mouse) or PBS. We monitored weight and survival over time (**Figure S4A**). We found that IFN-β treatment started one day after WT VACV infection at 2 × 10^5^ pfu or 2 × 10^6^ pfu successfully slowed down weight loss and protected mice from lethality (**Figure S4B-S4E**). Taken together, our results indicate that IFN-β production and signaling in the lungs are critical for host defense against vaccinia infection.

### Intranasal infection of VACVΔC7L results in the influx of dendritic cells (DCs), monocytes, neutrophils, CD8^+^, and CD4^+^ T cells into bronchoalveolar space of the infected lungs

To understand the reduced virulence of VACVΔC7L compared with WT VACV in the intranasal infection model, we performed immune cell analyses of bronchoalveolar lavage fluid (BAL) of WT VACV or VACVΔC7L-infected mice. The mice were infected either with VACV at 2 × 10^5^ pfu or with VACVΔC7L at 2 × 10^7^ pfu, or mock-infected with PBS. BAL was collected at 3 and 6 days post infection or PBS treatment. We chose to infect the mice at a lower pfu of WT VACV because of its high virulence in the intranasal infection model. We observed that Siglec F^+^CD11c^+^ lung resident AMs comprise the majority of CD45^+^ cells in the BAL in the PBS mock-infected mice. WT VACV infection resulted in the reduction of the absolute number of Siglec F^+^CD11c^+^ macrophages at day 6 post infection, with a mild increase of other myeloid cell populations in the BAL compared with mock-infected controls (**Figures 5A-5D; Figure S5A-S5D**). By contrast, VACVΔC7L infection caused a large influx of CD45^+^ myeloid cells, which included Ly6C^+^CD11b^+^ inflammatory monocytes, Ly6G^+^ neutrophils, and MHCII^+^CD11c^+^ DCs, into bronchoalveolar space at day 6 post infection (**Figures 5A-5D; Figure S5A-S5D**). DCs are important for presenting viral antigens to naïve T cells to generate antiviral T cells in the draining lymph nodes. The increased recruitment of DCs into the alveolar space positively correlates with the increased CD4^+^ and CD8^+^ T cells in the BAL at day 6 after VACVΔC7L infection (**Figures 5E-5G**). We assessed viral-specific CD8^+^ T cells responses by stimulating them with vaccinia dominant B8 epitope TSYKFESV and performing intracellular IFN-γ cytokine staining. SIINFEKL peptide, an irrelevant epitope from chicken ovalbumin, was used as a negative control. We found that VACVΔC7L infection resulted in the extravasation of viral-specific CD8^+^ T cells into the BAL (**Figures 5H-5I**). Taken together, these results indicate that VACVΔC7L infection leads to the recruitment of dendritic cells, monocytes, neutrophils, CD8^+^, and CD4^+^ T cells into the bronchoalveolar space of the infected lungs, whereas WT VACV infection has only a mild effect.

**Figure 5.**
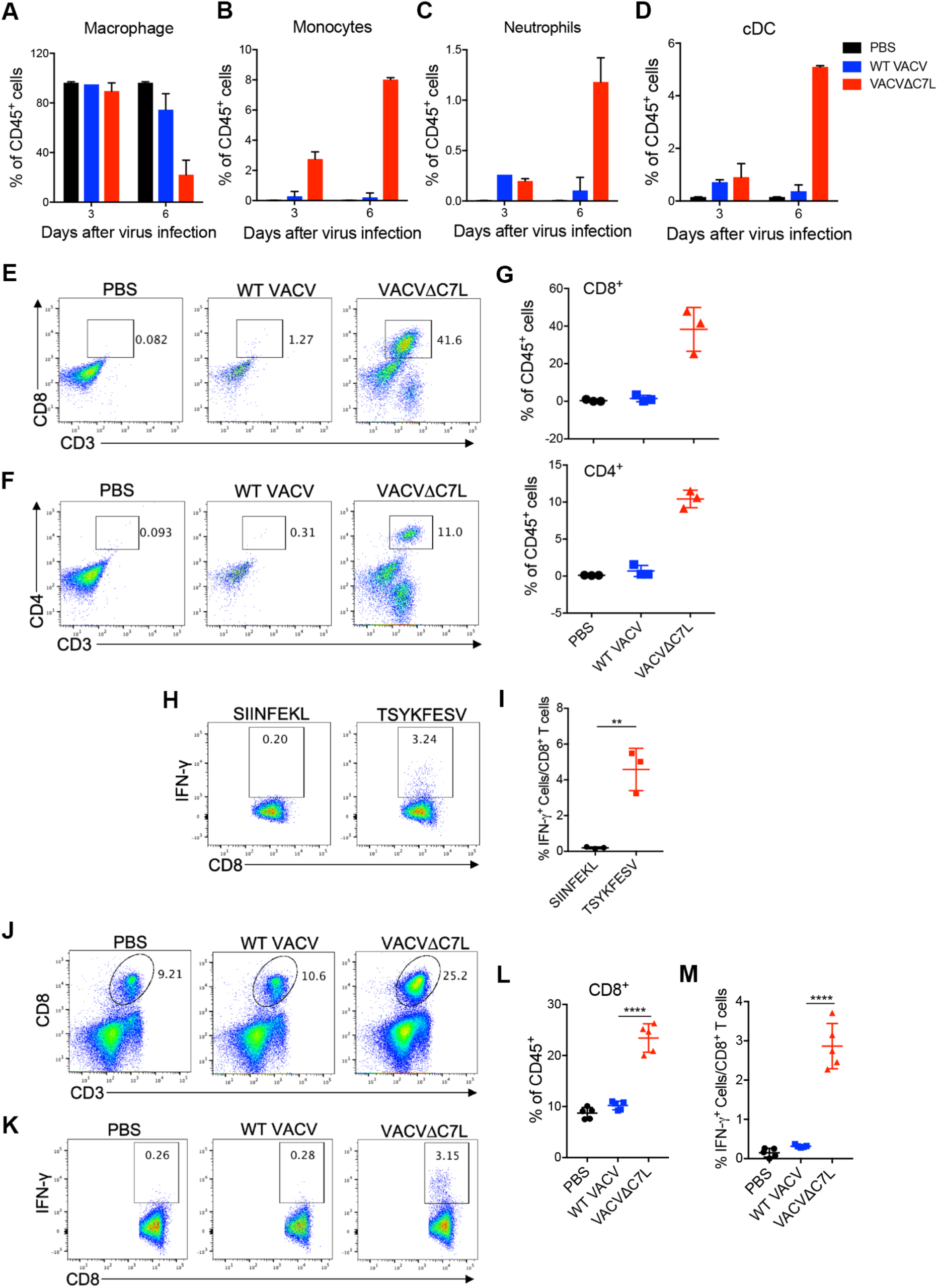
Intranasal infection of VACVΔC7L results in the recruitment of monocytes, DCs, neutrophils, CD8^+^, and CD4^+^ T cells into the infected lungs. (A-D) graphs of percentages of alveolar macrophages, monocytes, DCs, neutrophils out of CD45+ cells in the BAL of mice at day 3 and 6 post infection with either WT VACV or VACVΔC7L. A representative experiment is shown, repeated once. (E and F) Dot plots of CD8^+^ and CD4^+^ T cells in the BAL of mice at day 6 post infection with either WT VACV or VACVΔC7L. PBS was used as mock infection control. (G) Graphs showing percentages of CD8^+^ and CD4^+^ T cells out of CD45^+^ cells in the BAL of mice at day 6 post infection with WT VACV or VACVΔC7L. (H and I) Dot plots (H) and graph (I) showing B8 epitope (TSYKFESV)-specific IFN-γ^+^ CD8^+^ T cells in the BAL of mice at day 6 post infection with VACVΔC7L. SIINFEKL peptide was used as a negative control (n=3). (J and L) Dot plots (J) and graph (L) showing percentages of CD8^+^ T cells out of CD45^+^ cells in the lungs of mice at day 6 post infection with either WT VACV or VACVΔC7L. PBS was used as mock infection control (n=5). (K and M) Dot plots (K) and graph (M) showing B8-specific IFN-γ^+^ CD8^+^ T in the lungs of mice at day 6 post infection with either WT VACV or VACVΔC7L. PBS was used as mock infection control (n=5). A representative experiment is shown, repeated once.

### Intranasal infection of VACVΔC7L generates more viral-specific activated CD8^+^ T cells in the infected lungs compared with WT VACV

We analyzed CD8^+^ T cells in lungs at day 5 post infection with either WT VACV or VACVΔC7L. While WT VACV infection had a limited effect on the percentages of CD8^+^ T cells out of CD45^+^ cells in the lungs, VACVΔC7L infection strongly boosted CD8^+^ T cells infiltration into lungs (**Figures 5J and 5L**). We also found that VACVΔC7L infection resulted in higher percentages of vaccinia B8-specific CD8^+^ T cells in lungs compared with WT VACV (**Figures 5K and 5M**). These results suggest that host innate immunity, including type I IFN induced by VACVΔC7L might facilitate the generation of viral-specific CD8^+^ T cell responses.

### T, B, NK, and alveolar macrophages are dispensible for host restriction of VACVΔC7L infection in an intranasal infection model

VACVΔC7L infection at 2 × 10^7^ pfu in RAG1-deficient mice, which lack T and B cells, resulted in only mild weight loss, and all of the mice recovered their weight around day 10 and 11 post infection and survived (**Figures 6A-6B**). Antibody depletion of NK cells did not affect VACVΔC7L-induced weight loss and did not enhance mortality (**Figures 6C-6D**). Furthermore, intranasal application of liposomal clodronate resulted in depletion of alveolar macrophages, However, this treatment did not exacerbate VACVΔC7L-induced weight loss. These results indicate that T, B, NK, and alveolar macrophages are not important for controlling VACVΔC7L pulmonary infection in this intranasal infection model.

**Figure 6.**
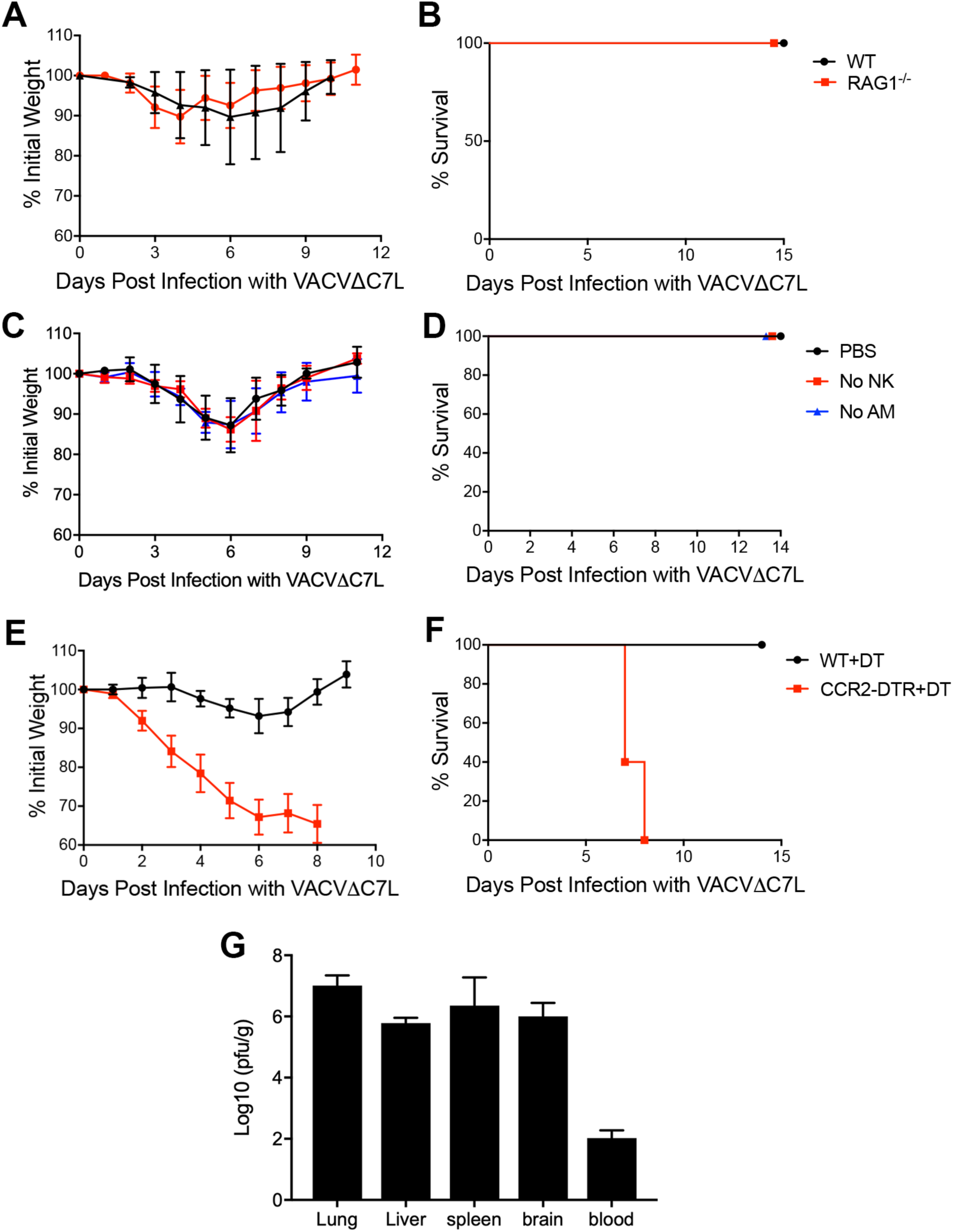
CCR2^+^ inflammatory monocytes plays important roles in restricting VACVΔC7L infection in the lungs and in preventing systemic dissemination. (A) shown are the percentages of initial weight over days post intranasal infection with VACVΔC7L at 2 × 10^7^ pfu in RAG1^−/−^ and age-matched WT C57BL/6J mice. (B) Kaplan-Meier survival curve of RAG1^−/−^ and age-matched WT C57BL/6J mice infection with VACVΔC7L at 2 × 10^7^ pfu (n=5 in each group). (C) shown are the percentages of initial weight over days post intranasal infection with VACVΔC7L at 2 × 10^7^ pfu in WT C57BL/6J mice treated with anti-NK antibody or with liposomal clodronate delivered intranasally. (D) Kaplan-Meier survival curve of WT C57BL/6J mice treated with anti-NK antibody or with liposomal clodronate infected with VACVΔC7L at 2 × 10^7^ pfu (n=5 in each group). A representative experiment is shown, repeated once. (E) shown are the percentages of initial weight over days post intranasal infection with VACVΔC7L at 2 × 10^7^ pfu in CCR2-DTR and age-matched WT C57BL/6J mice treated with DT. (F) Kaplan-Meier survival curve of CCR2-DTR and age-matched WT C57BL/6J mice treated with DT infected with VACVΔC7L at 2 × 10^7^ pfu (n=5 in each group). A representative experiment is shown, repeated once. (G) Titers of VACVΔC7L in the lungs, livers, spleens, blood, and brains of CCR2-DTR mice treated with DT at day 7 or 8 post intranasal infection with VACVΔC7L at a dose of 2 × 10^7^ pfu. Data are represented as mean ± SEM (n=3-5).

### CCR2^+^ inflammatory monocytes contribute to host restriction of VACVΔC7L infection in the lungs

Our immune cell profiling of BAL showed that Ly6C^+^CD11b^+^ inflammatory monocytes are the major myeloid cell population extravasated into the BAL at day 3 and 6 post VACVΔC7L infection (**Figures 5B and S3B**). To examine the function of CCR2^+^Ly6C^hi^ inflammatory monocytes in host defense against VACVΔC7L infection, we used CCR2-DTR mice to transiently deplete CCR2^+^ monocytes prior to VACVΔC7L infection by administering diphtheria toxin (DT) intraperitoneally (Hohl et al., 2009). We found that CCR2-DTR mice treated with DT were much more susceptible to VACVΔC7L infection compared with WT mice treated with DT, with more rapid weight loss and 100% mortality at day 7 or 8 post infection (**Figures 6E-6F**). We determined viral titers in various organs of the mice that were euthanized because of their loss of more than 30% of original weight. We found that depletion of CCR2^+^ monocytes resulted in viremia and systemic dissemination. These results demonstrate that CCR2^+^ inflammatory monocytes play an important role in controlling vaccinia infection in the lungs and in preventing systemic dissemination.

### CCR2^+^ inflammatory monocytes differentiate into interstital macrophages (IMs), DCs in the lungs upon VACVΔC7L infection

To track the CCR2^+^ monocytes after intranasal viral infection, we used CCR2-GFP reporter mice in which enhanced GFP is expressed in CCR2^+^ cells under the control of CCR2 promoter (Hohl et al., 2009). Intranasal infection of WT mice with VACVΔC7L leads to a marked increase of GFP^+^ Ly6C^+^ inflammatory monocytes, IMs (especially Lyve1^−^ IMs), and a modest increase of CD11c^+^ DCs in the infected lungs at 3 days post infection (**Figure 7A**). Using single-cell RNA sequencing analysis, it has been recently shown that CCR2^+^Ly6C^+^ monocytes can differentiate into two distinct IM populations, Lyve1^lo^MHCII^hi^ and Lyve1^hi^MHCII^lo^, in the lungs (Chakarov et al., 2019). Whether the two IM popuations have distinct roles in antiviral activities need to be explored in future studies. Taken together, our results indicate that intranasal infection of VACVΔC7L leads to the recruitment of CCR2^+^ monocytes into the lungs, which can further differentiate into IMs and DCs under the influence of an inflammatory milieu in the infected lungs.

**Figure 7.**
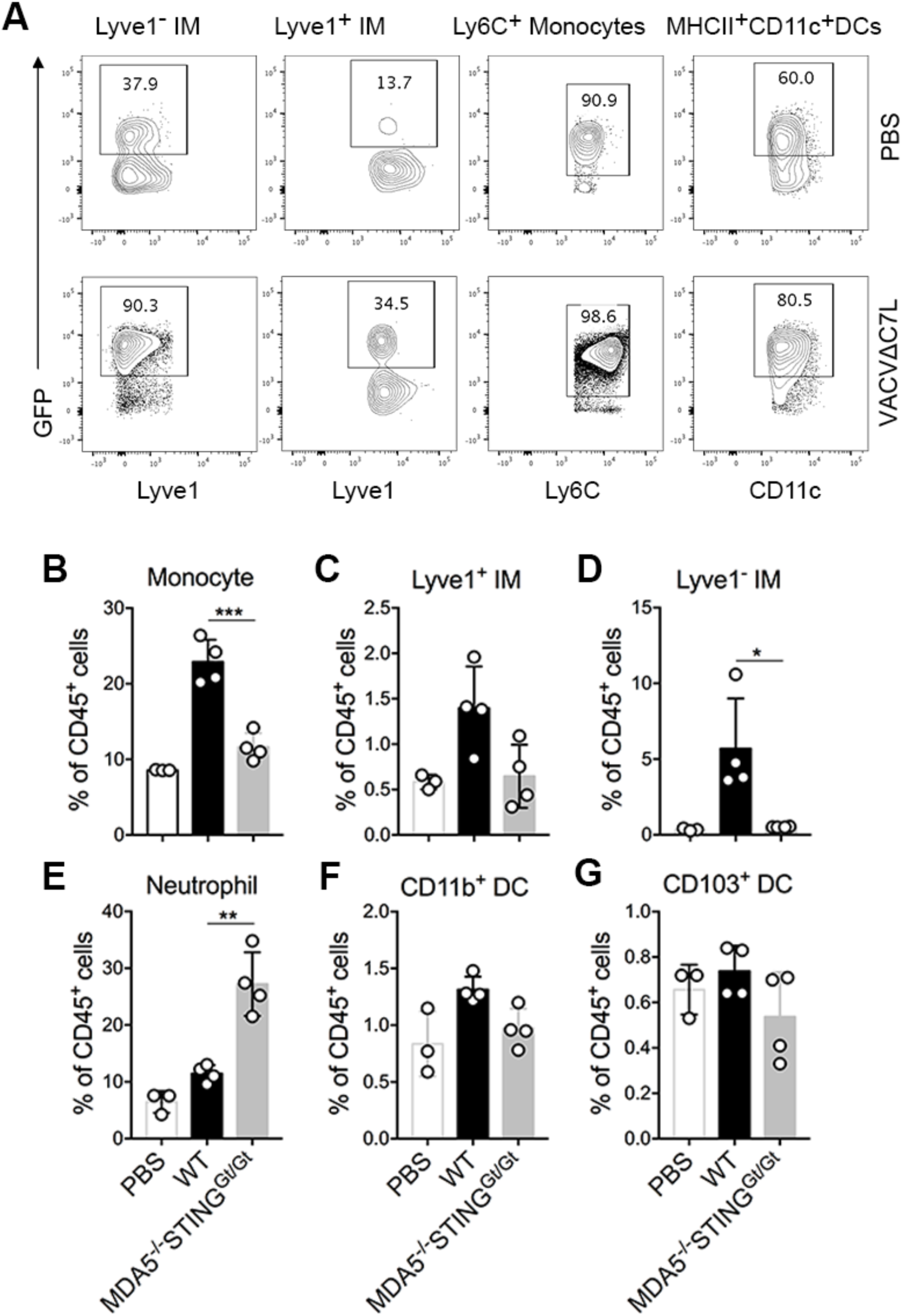
CCR2^+^ inflammatory monocytes differentiate into interstital macrophages (IMs), DCs in the lungs upon VACVΔC7L infection. (A) dot plots showing an increase of GFP^+^ Lyve1^−^ IMs, GFP^+^Lyve1^+^ IMs, and GFP^+^Ly6C^+^ monocytes and GFP^+^MHCII^+^CD11C^+^ DCs in the lungs of CCR2-GFP mice at day 3 post infection with VACVΔC7L compared with PBS-mock infected mice. (B-G) graphs showing percentages of Ly6C^+^ monocytes, Lyve1^+^ IMs, Lyve1^−^ IMs, Ly6G^+^ neutrophils, CD11b^+^ DCs, and CD103^+^ DCs out of CD45^+^ cells in the lungs of WT and MDA5^−/−^ STING^Gt/Gt^ mice at day 3 post VACVΔC7L infection. Data are represented as mean ± SEM (n=3-4). PBS mock infection control was performed in WT mice.

The MDA5^−/−^STING^Gt/Gt^ mice are highly susceptible to VACVΔC7L infection (**Figures 1G-1H and 1J**). The lung AECIIs from MDA5^−/−^STING^Gt/Gt^ mice fail to induce the expression of IFNB and inflammatory cytokine and chemokine genes upon VACVΔC7L infection (**Figures 2H and 2I**). We hypothesized that the lack of induction of innate immunity in the lung epithelium of MDA5^−/−^STING^Gt/Gt^ mice would result in the failure of recruiting CCR2^+^ monocytes. To test that, we performed intranasal infection of VACVΔC7L in WT and MDA5^−/−^STING^Gt/Gt^ mice. PBS was used as a mock infection control in WT mice. Lungs were collected at day 3 post infection and we analyzed myeloid cell populations. We found that intranasal infection of VACVΔC7L resulted in the recruitment of Ly6C^+^ inflammatory monocytes, and the generation of Lyve1^+^ and Lyve1^−^ IMs in WT mice (**Figures 7B-7D**). However, VACVΔC7L induced monocytes and Lyve1^−^ IMs were markedly reduced in MDA5^−/−^STING^Gt/Gt^ mice (**Figures 7B-7D**). This indicates that innate immunity mediated by the cytosolic DNA and dsRNA-sensing pathways most likely in the lung AECs is critical for the recruitment of CCR2^+^ monocytes into the lungs as well as their differentiation into Lyve1^−^ IMs.

## DISCUSSION

In this study, we established an acute pulmonary DNA virus infection model with an attenuated but replication-competent mutant VACV with the deletion of a host range protein encoded by the C7L gene (VACVΔC7L). WT VACV is virulent in C57BL/6J mice. The LD_50_ is around 2 × 10^5^ pfu given intranasally. By contrast, VACVΔC7L is non-virulent given at 2 × 10^7^ pfu. Using genetic knock-out mice, or antibody or chemical depletion methods, we demonstrated that T, B, NK, and alveolar macrophages are dispensable in this acute viral infection model. On the contrary, the innate immune system consisting of lung AECII and CCR2^+^ monocytes play important roles combating against acute high-dose infection with this replicative DNA virus. Our results also demonstrate that both the cytosolic dsRNA-sensing pathway mediated by MDA5 and the cytosolic DNA-sensing pathway mediated by cGAS/STING play important roles in host defense against VACVΔC7L infection. Our results support a model in which VACVΔC7L infection triggers IFN-β and CCL2 production from lung AECII, which strengthens an antiviral state through activating the IFN-β/IFNAR/STAT2 pathway, as well as recruiting CCR2^+^ inflammatory monocytes through the CCL2/CCR2 axis. In the infected lungs and under the influence of various cytokines and chemokines, CCR2^+^ inflammatory monocytes then further differentiate into interstitial macrophages and dendritic cells to fortify host immunity.

Vaccinia virus encodes many immunomodulatory genes to evade the host immune system (Brady and Bowie, 2014; Seet et al., 2003). In this study, we focused on the host-range protein C7 and its inhibitory effect on IFN production. Vaccinia C7 was discovered as a host range protein that allows vaccinia replication in human cells (Perkus et al., 1990). It is functionally equivalent to another vaccinia host range protein K1 and therefore deletion of both C7L and K1L gene from the vaccinia genome renders the virus replication-incompetent in certain human cells (Perkus et al., 1990). One of the myxoma homologs of C7 encoded by the M62R has been shown to interact with host factor SAMD9 protein in human cells (Liu et al., 2011). Through an unbiased genome-wide siRNA screen in human cells, SAMD9 and WDR6 were identified as host restriction factors for vaccinia virus lacking both the C7L and K1L genes (Sivan et al., 2015). C7 can also bind to SAMD9L in mouse, and VACV with both C7L and K1L deletion is highly attenuated in an intranasal infection model, but it gains virulence in SAMD9L^−/−^ mice (Meng et al., 2018). The human SAMD9 and murine SAMD9L genes are ISGs (Meng et al., 2018; Tanaka et al., 2010). Through screening a library of more than 350 human ISGs, overexpression of transcription factor IRF1 was identified to be able to suppress the replication of mutant vaccinia with deletions of C7L and K1L (Meng et al., 2012). However, VACVΔC7L or VACVΔK1L replication was insensitive to IRF1 overexpression, suggesting that both C7 and K1 antagonize IRF1-mediated inhibitory effects (Meng et al., 2012).

Using a dual-luciferase reporter assay, we found that overexpression of C7 attenuates STING-, TBK-, TRIF, MAVS-, TLR3/poly (I:C) and IRF3-induced IFNB promoter activation in HEK293-T cells. Co-immunoprecipitation studies revealed that C7 interacts with IRF3. Although neither WT VACV nor VACVΔC7L infection of myeloid cells including AMs and BMDCs induces IFNB gene expression or IFN-β protein production, MVAΔC7L infection of BMDCs induces higher levels of IFNB gene expression and IFN-β secretion compared with MVA. Furthermore, MVAΔC7L infection induces higher levels of phosphorylation of IRF3 compared with MVA, indicating that C7 might prevent IRF3 phosphorylation.

More strikingly, VACVΔC7L is attenuated by more than 100-fold compared with WT VACV in WT C57BL/6J mice in an intranasal infection model. We attribute the attenuation of VACVΔC7L to the following key factors: (i) VACVΔC7L infection of murine primary lung alveolar epithelial cells results in the induction of Ifnb and ISG gene expression in a MDA5/STING-dependent manner; (ii) intranasal infection of VACVΔC7L leads to the recruitment of DCs, monocytes, neutrophils, CD8^+^, and CD4^+^ T cells into the BAL and lung parenchyma; and (iii) intranasal infection of VACVΔC7L leads to the recruitment of CCR2^+^ inflammatory monocytes into the lung parenchyma and their differentiation into interstitial macrophages. These results established that the host range factor vaccinia C7 is a key virulence factor and VACVΔC7L infection triggers innate immunity in the lungs that results in restriction of viral replication and spead.

Murine lung epithelial cells and alveolar macrophages provide the front-line defense against invading viral pathogens during pulmonary infection. In the lower respiratory tract, the murine lung epithelial cell lining is comprised of two main cell types, type I and type II alveolar epithelial cells (AECIs and AECIIs). AECIIs have been shown to be the major targets of influenza virus infection and contribute to the innate immune defense against viral pathogens both in vitro and in vivo (Galani et al., 2017; Stegemann-Koniszewski et al., 2016; Weinheimer et al., 2012; Yu et al., 2011). It was recently shown that AECIIs sorted from the lungs of mice infected with influenza virus (IAV) up-regulate the expression of many antiviral factors and immune mediators. This correlates with the ability of the virus to recruit immune cells into the BAL of the infected lungs (Stegemann-Koniszewski et al., 2016).

The role of lung AECIIs in host defense against VACV infection has not been demonstrated previously. In this study, we isolated lung AECIIs by using an anti-EpCAM antibody and then cultured these cells in the presence of keratinocyte growth factor on matrigel-coated plates. We found that VACVΔC7L infection of AECIIs induces the expression of Ifnb and ISGs, whereas WT VACV fails to do so. Using lung AECII isolated from MDA5 and STING-double deficient mice, we found that VACVΔC7L-induced Ifnb and ISG gene expression is dependent on the cytosolic nucleic acid-sensing pathways.

Alveolar macrophages (AMs) have been shown to play some roles in host defense against vaccinia infection by using *Csf2-/-* mice, which lack AMs, or by using liposomal clodronate to delete AMs, when the mice were infected at 10^4^ to 10^5^ pfu (Rivera et al., 2007; Schneider et al., 2014). The protective roles of AMs were also observed with intratracheal infection with PR8 influenza virus (Schneider et al., 2014). It has been postulated that AMs play a protective role in the lungs by removing dead cells and eosinophilic surfactant material aggravated by viral infection. In this study, depletion of AMs by liposomal clodronate did not affect the weight loss or mortality induced by intranasal infection of VACVΔC7L at 2 × 10^7^ pfu. We also observed that neither VACVΔC7L nor WT VACV infection of AMs induces IFN or proinflammatory cytokine/chemokine production. Therefore, unlike lung AECIIs, AMs do not seem to be important in host defense against VACVΔC7L intranasal infection in WT mice.

To understand which cell population(s) are the major producer(s) of IFN-β after intranasal infection with VACVΔC7L, we used IFNβ-yellow fluorescent protein (YFP) knockin mice to map the IFN-β producing cells in the lungs after VACVΔC7L infection (Scheu et al., 2008). FACS results show that the majority of YFP^+^ cells in the lungs at 1 day post VACVΔC7L intranasal infection are CD45^−^CD31^−^T1a^−^EpCAM^+^, which are consistent with lung AECIIs. Immunohistochemistry (IHC) of lungs collected at 1 day post VACVΔC7L infection showed that among the AECII cells, which are positive for surfactant protein C (SPC), some are also positive for IFNβ-YFP as detected by anti-GFP antibody. These results are congruent with our *in vitro* results that VACVΔC7L infection of purified lung AECs induces IFNB gene expression and IFN-β protein secretion.

The type I IFN receptor and JAK/STAT pathway are critical for host resistance to viral infection. The IFNAR1-deficient mice are more susceptible to WT VACV infection compared with WT mice (Muller et al., 1994; van den Broek et al., 1995). Because STAT2 acts downstream of IFNAR1, it is expected that STAT2^−/−^ mice may also be more susceptible to vaccinia infection. The difference of virulence of VACVΔC7L in WT and STAT2^−/−^ or IFNAR1^−/−^ mice is striking. Whereas VACVΔC7L at 2 × 10^7^ pfu causes only transient weight loss but no lethality in WT mice, its LD50 in STAT2^−/−^ or IFNAR1^−/−^ mice is around 1000 pfu.

Our bone marrow chimera results indicate that the type I IFN feedback loop in the non-hematopoietic cell population(s) is most important for mice survival after VACVΔC7L intranasal infection, although IFNAR1 signaling in hematopoietic cell population(s) also contributes to host defense. Using mice with specific deletion of Ifnar1 in lung AECIIs (Ifnar1^fl/fl^-Sftpc^cre-ERT2^), we found that type I IFN signaling on lung AECII is critical for host defense against VACVΔC7L intranasal infection. Taken together, these results demonstrate that lung AECIIs produce IFN-β in response to VACvΔC7L infection, which in turn directly stimulates IFNAR1 on AECIIs to restrict viral replication and spread. In addition, they produce chemokines to recruit hematopoietic innate immune cells to the infected tissue for boosting antiviral immunity.

Monocytes egress from the bone marrow to the blood circulation in response to infection and they also migrate to the infected or inflamed tissue in a CCR2-dependent manner where they further differentiate into other cell types, including inflammatory DCs and macrophages. CCR2^+^ monocytes have been shown to be important for antiviral immunity in a mouse model of intravaginal infection with herpes simplex virus 2 (HSV-2). CCR2^−/−^ mice are more susceptible to HSV-2 infection with worsening clinical symptoms, increased mortality, and higher viral titers in vaginal wash (Iijima et al., 2011). By contrast, in an intranasal infection model of influenza virus, CCR2^+^ monocytes contribute to the influenza-induced lung immunopathology and death (Lin et al., 2008). To address the role of CCR2^+^ monocytes in host defense against intranasal infection with VACVΔC7L, we used CCR2-DTR mice, in which CCR2^+^ monocytes can be transiently depleted by intraperitoneal delivery of diphtheria toxin (DT) (Hohl et al., 2009). We found that depletion of CCR2^+^ inflammatory monocytes renders the mice susceptible to VACVΔC7L infection. All of the infected CCR2-DTR mice died after DT treatment with viremia and systemic dissemination of the virus. To understand the role of CCR2^+^ monocytes in host defense against vaccinia infection, we used CCR2-GFP reporter mice to track the CCR2^+^ monocytes after intranasal infection with VACVΔC7L. Our results show that VACVΔC7L infection in WT mice causes the recruitment of CCR2^+^ monocytes into the infected lungs and their differentiation into Lyve1^−^ IMs. These effects were lost in VACVΔC7L-infected MDA5^−/−^ STING^Gt/Gt^ mice.

In conclusion, using an attenuated vaccinia virus (VACVΔC7L), we established an intranasal DNA virus infection model in which adaptive immunity is dispensable for host defense against acute infection. This allows us to focus on host innate immunity in restricting viral infection in the lungs. Our results highlight the cross-talk between lung AECs with CCR2^+^ monocytes in the control of acute pulmonary viral infection.

## ACKNOWLEDGEMENTS

We thank the Flow Cytometry Core Facility and Molecular Cytology Core Facility at the Sloan Kettering Institute. We thank Stewart Shuman, Eric Pamer, Jedd Wolchok, and Taha Merghoub for helpful discussions. We thank Joan Libermann-Smith for editing. This work was supported that NIH grant K-08 AI073736 (L.D.), R56AI095692 (L.D.), Lucille Castori Center for Microbes, Inflammation & Cancer seed grant (L.D.), the Society of Memorial Sloan Kettering (MSK) research grant (L.D.), MSK Technology Development Fund (L.D.), Sponsored Research Award from IMVAQ Therapeutics. LD is the recipient of a Physician Scientist Career Development Award from the Dermatology Foundation, a research scholar from American Skin Association. She is the recipient of a career development award from Parker Institute for Cancer Immunotherapy. This research was also funded in part through the NIH/NCI Cancer Center Support Grant P30 CA008748.

## AUTHOR CONTRIBUTIONS

Author contributions: L.D. and N.Y. designed and performed the experiments, analyzed the data, and prepared the manuscript. P.D. and Y.W. assisted in some experiments, analyzed the data. J.L. assisted in some experiments, analyzed the data, and assisted in manuscript preparation. C.M.R. assisted in experimental design, data interpretation, and manuscript preparation. Memorial Sloan Kettering Cancer Center filed a patent application for the use of recombinant MVAΔC7L or VACVΔC7L as monotherapy or in combination with immune checkpoint blockade for solid tumors and vaccine applications.

## Materials and Methods

### Mice

Female C57BL/6J mice between 6 and 8 weeks of age were purchased from the Jackson Laboratory and were used for the preparation of bone marrow-derived dendritic cells and for intranasal infection experiments. IFNb/YFP reporter mouse, cGAS^−/−^, STAT2^−/−^, IFNAR1^−/−^, Sftpc-CreER^T2^, Ifnar^fl^, mice were purchased from the Jackson Laboratory. STING^Gt/Gt^ mice were generated in the laboratory of Russell Vance (University of California, Berkeley). MDA5^−/−^ mice were generated in Marco Colonna’s laboratory (Washington University). MDA5^−/−^STING ^Gt/Gt^ and Ifnar1^fl/fl^-Sftpc^cre-ERT2^ mice were breeded in our lab. CCR2-GFP and CCR2-DTR mice were provided by Eric Pamer (Memorial Sloan Kettering Cancer Center). These mice were maintained in the animal facility at the Sloan Kettering Institute. All procedures were performed in strict accordance with the recommendations in the Guide for the Care and Use of Laboratory Animals of the National Institute of Health. The protocol was approved by the Committee on the Ethics of Animal Experiments of Sloan-Kettering Cancer Institute.

### Intranasal infection of WT VACV or VACVΔC7L in mice

5-10 WT mice in each group were anesthetized and infected intranasally with increasing doses of WT VACV or VACVΔC7L at indicated pfu, inoculated to both nostrils in 20 µl each. Mice were monitored and weight daily. Mice that had lost over 30% of initial weight were be euthanized. Kaplan-Meier survival curves were determined.

### Cytokine production assays

For in vivo experiments, 1 ml of bronchoalveolar lavage was used for cytokine measurements. For in vitro experiments, cell supernatant was collected for analysis. Most cytokines were measured using commercial mouse ELISA kits. IFN-β was measured by ELISA (PDL) and CCL4 and CCL5 were measured by ELISA (R&D). The Luminex assay was performed using the Cytokine Mouse Magnetic 20-Plex Panel (ThermoFisher).

### Flow cytometry

To analyze cell populations in the BALF and lung, lungs were digested with Collagenase D (2mg/ml) and DNase I (100 µg/ml) for 45 mins at 37°C. Single Cell suspensions were blocked Anti-CD16/CD32 antibody and stained with antibodies for 30 mins on ice. LIVE/DEAD™ Fixable Aqua Stain (ThermoFisher) was used to stain dead cells. Fluorescently conjugated antibodies, including anti-CD45.2 (clone 104), anti-Ly6G (clone 1A8), anti-Ly6C (clone HK1.4), anti-CD11c (clone N418), anti-CD11b (clone M1/70), anti-MHC II (clone M5/114.15.2), anti-CD31 (clone MEC13.3), anti-EpCAM (clone G8.8), anti-CD104 (clone 346-11A), anti-CD64 (clone X54-5/7.1) anti-CD3ε (clone 145-2C11), anti-CD4 (clone RM4-5), anti-CD8 (clone 53-6.7) and anti-IFN-γ (clone XMG1.2) were from BioLegend. Anti-CD45 (clone 30-F11), anti-Siglec F (clone E50-2440) were from BD biosciences. Anti-MERTK (clone DS5MMER) was from ThermoFisher. For intracellular cytokine staining, cell suspensions were incubated with 5 µg/ml peptide (B8R 20-27 or OVA 257-264) and Brefeldin A (0.1%) for 4 hours at 37°C prior to all staining, treated with BD Cytofix/Cytoperm™ kit for staining. Cells were analyzed on the BD LSR II flow cytometer. Data were analyzed with FlowJo software (version 10.5.3).

### Tissue fixation and immunostaining

Lungs were fixed with 4% paraformaldehyde for overnight at 4°C. Lungs were embedded in O.C.T and cryosections (10 µm) were used for immunofluorescent (IF) analysis. Tissue sections were permeabilized with 0.5% Triton X-100 in PBS for 5 min. Then blocked in 5% goat serum (Sigma), 3% bovine serum albumin (Fisher) and 0.1% Triton X-100 for 1 hr at room temperature. Primary antibodies were incubated overnight at 4°C at the indicated dilutions: chicken anti-GFP (1:1000, Abcam), rabbit anti-SP-C (1:1000, Millipore). Alexa Fluor-coupled secondary antibodies (1:1000, Invitrogen) were incubated at room temperature for 60 min. After antibody staining, sections were embedded in ProLong Gold Antifade Mountant (ThermoFisher). Images were acquired using a confocal microscope (Leica TCS SP8). All the images were further processed with Image J software.

### Tamoxifen and Diphtheria Toxin Administration

Diphtheria toxin was obtained from Sigma, reconstituted at 1 mg/ml in PBS, and frozen at −80°C. Mice received 10 ng/g DT via the i.p. route in 0.2–0.3 ml PBS. Tamoxifen (Sigma) was a 40 mg/ml stock solution in corn oil (Sigma) and given 4 mg via intraperitoneal (IP) injection × 4-5 doses.

### Viruses and Cell lines

The WR strain of vaccinia virus (VACV) was propagated and virus titers were determined on BSC40 (African green monkey kidney cells) monolayers at 37°C. MVA virus was kindly provided by Gerd Sutter (University of Munich), and propagated in BHK-21 (baby hamster kidney cell, ATCC CCL-10) cells. The viruses were purified through a 36% sucrose cushion. Heat-iMVA was generated by incubating purified MVA virus at 55 °C for 1 hour. Sendai virus (SeV; Cantell strain) was obtained from Charles River Laboratories. BSC40, HEK293T and RAW264.7 were cultured in Dulbecco’s modified Eagle’s medium supplemented with 10% fetal bovine serum (FBS), 2 mM L-glutamine and 1% penicillin-streptomycin. BHK-21 were cultured in Eagle’s Minimal Essential Medium (Eagle’s MEM, can be purchased from Life Technologies, Cat# 11095-080) containing 10% FBS, and 1% penicillin-streptomycin. For THP-1 differentiation into macrophages, they were treated with PMA (10 ng/ml) for 72 h.

To culture primary murine AEC2, the lungs from mice (6-8 weeks) were perfused via the right ventricle with 10 ml PBS, then inflated with a 1.5 ml mixture of 1 ml low melting agarose (1% w/v) and 500 µl dispase (Corning). The lung lobes were gently minced into small pieces in a conical tube containing 3ml of PBS, 1U/mL of dispase (Roche), and 100U/ml DNase I (Sigma) followed by rotating incubation for 45 min at 37°C. The cells were filtered through 40 µm mesh and for further staining against antibodies for mouse flow cytometry: pan CD45-APC, CD31-APC, FITC-CD104 and EpCAM-APC-Cy7 (BioLegend). LIVE/DEAD™ Fixable Aqua Stain (ThermoFisher) was used to eliminate dead cells. Cell sorting was performed with a FACS Aria II (BD Biosciences), and data were analyzed with FlowJo software (Tree Star, Inc.). AECII progenitors cells were plated into a Matrigel (Corning, 354230) pre-coated TC plate and cultured with Small Airway Epithelial Cell Growth Medium (Lonza) supplemented with charcoal-stripped 5% FBS, 10 ng/ml keratinocyte growth factor (PeproTech, 100-19), 10 µM Rock inhibitor (Selleck Chemicals, S1049), and 1% P/S at 37°C in a 5% CO2 incubator for the first 2 days, and then replaced with the same media but without Rock inhibitor for the next 4 to 5 days.

### Multistep growth curve of WT VACV and VACVΔC7L

AEC2 cells were infected with WT VACV or VACVΔC7L at a MOI of 0.05. The cells were then scraped into the medium and collected at indicated times. After three cycles of freeze-thaw and subsequent sonication, viral titers in the collected samples were determined by plaque assay on BSC40 cells.

### Plasmid Construction

IFN-β reporter plasmid (pIFN-β-luc) and ISRE reporter plasmid (p-ISRE-luc) were provided by Michaela Gack (University of Chicago). STING, TBK1, IRF3 were provided by Tom Maniatis (University of Columbia). IRF3-5D were provided by Rongtuan Lin (McGill University). MAVS, TLR3, TRIF plasmids were purchased from Addgene. VACV C7L was amplified by PCR from VACV WR genome and subcloned into pcDNA3.1 and pQCXIP.

### Dual Luciferase Reporter assay

Luciferase activities were measured using the Dual Luciferase Reporter Assay system according to the manufacturer’s instructions (Promega). Briefly, expression plasmids including a firefly luciferase reporter construct, a *Renilla* luciferase reporter construct, as well as other expression constructs were transfected into HEK293T cells. 24 h post transfection, cells were collected and lysed. The relative luciferase activity was expressed as arbitrary units by normalizing firefly luciferase activity under IFNB promoter to Renilla luciferase activity from a control plasmid pRL-TK.

### Construction of retrovirus expressing vaccinia C7L

HEK293T cells were passaged into 6-well plate. Next day, cells were transfected with three plasmids-VSVG, gag/pol and pQCXIP-C7 or pQCXIP with lipofectamine 2000. After 2 days, cell supernatants were collected and filted through 0.45 µm filter and stored in −80 °C.

### Generation of RAW264.7 and THP-1 cell line stably expressing vaccinia C7L

Cells were passaged into 6-well plate. Next day, cells were infected with retrovirus expressing C7L or control virus at MOI 5. After 2 days, culture medium was replaced with medium containing puromycin. After one week, survival cells are the cells stably expressing C7L and verified by Western blot analysis using anti-C7 antibody.

### Generation of recombinant VACVΔC7L virus

BSC40 cells were passaged into 6-well plate. Next day, cells were infected with WT vaccinia virus WR strain at MOI 0.2. After 1-2 h, cells were transfected with pC7-GFP or pC7-mCherry with lipofectamine 2000. Both GFP and mCherry expression were under vaccinia synthesic early and late promoter (pSE/L). After 2 days, cells were collected and underwent three cycles of freeze/thaw. To select pure recombinant VACVΔC7L, BSC40 cells were infected with virus mix above, then select single plaques based on the GFP or mCherry expression under the microscope. After several rounds plaque purification, pure recombinant VACVΔC7L-GFP and VACVΔC7L-mCherry were obtained. PCR analyses and sequencing were performed to make sure that the C7L gene was deleted from the VACV genome.

### Generation of recombinant MVAΔC7L virus

BHK21 cells were passaged into 6-well plate. Next day, cells were infected with MVA at MOI 0.2. After 1-2 h, cells were transfected with pC7-GFP with lipofectamine 2000. After 2 days, cells were collected and underwent three cycles of freeze/thaw. BHK21 cells were infected with virus stock collected above, then select plaques based on the GFP expression under the microscope. After several rounds of selection, all plaques were GFP positive. GFP-positive MVAΔC7L clones were amplified and the deletion of the C7L gene was confirmed by PCR analysis.

### Generation of bone marrow-derived dendritic cells (BMDCs)

The bone marrow cells from the tibia and femur of mice were collected by first removing muscles from the bones, and then flushing the cells out using 0.5 cc U-100 insulin syringes (Becton Dickinson) with RPMI with 10% FCS. After centrifugation, cells were re-suspended in ACK Lysing Buffer (Lonza) for red blood cells lysis by incubating the cells on ice for 1-3 min. Cells were then collected, re-suspended in fresh medium, and filtered through a 40-µm cell strainer (BD Biosciences). The number of cells was counted. For the generation of GM-CSF-BMDCs, the bone marrow cells (5 million cells in each 15 cm cell culture dish) were cultured in CM in the presence of GM-CSF (30 ng/ml, produced by the Monoclonal Antibody Core facility at the Sloan Kettering Institute) for 10-12 days. CM is RPMI 1640 medium supplemented with 10% fetal bovine serum (FBS), 100 Units/ml penicillin, 100 µg/ml streptomycin, 0.1mM essential and nonessential amino acids, 2 mM L-glutamine, 1 mM sodium pyruvate, and 10 mM HEPES buffer. Cells were fed every 2 days with fresh medium.

### Western Blot Analysis

BMDCs (1 × 10^6^) from WT and KO mice were infected with MVA or MVAΔC7L at a MOI (multiplicity of infection) of 10. Whole-cell lysates were prepared. Equal amounts of proteins were subjected to sodium dodecyl sulfate-polyacrylamide gel electrophoresis and the polypeptides were transferred to a nitrocellulose membrane. Phosphorylation of IRF3, and IRF3 were determined using respective antibodies (Cell Signaling). Anti-C7 antibody was used to determine C7 expression by MVA. Anti-glyceraldehyde-3-phosphate dehydrogenase (GADPH) or anti-β-actin antibodies (Cell Signaling) were used as loading controls.

### Co-immunoprecipitation

HEK293T cells were passaged into 10 cm plates. Next day, cells were transfected with Flag-IRF3 together with pcDNA3.1-C7. After two days, cells were lysed in Pierce IP lysis buffer on ice for 30 min. C7 antibody was added into cell lysis to final concentration 1 µg/ml. Then incubate at 4 °C overnight. Next day, protein A-agarose was added and incubate at 4 °C for 2 h. Wash agarose with IP lysis buffer for three times. Lastly, the proteins were denatured at 98 °C 5 min before loading on a SDS-PAGE.

### Quantitative real-time PCR

Total RNA was obtained from cultured cells with TRIzol reagent (Invitrogen). Cellular RNAs were reverse-transcribed and amplified by PCR using the Verso cDNA synthesis kit (Thermo Fisher) and SYBR^TM^ Green Master Mix (Thermo Fisher). Cellular RNAs were normalized to GAPDH levels. All assays were performed on an ABI 7500 system and analyzed with ABI 7500 SDS software v.1.3. The primers used are as follows:

> mIFNB forward 5’-TGGAGATGACGGAGAAGATG-3’,
>
> mIFNB reverse 5’-TTGGATGGCAAAGGCAGT-3’,
>
> mCCL4 forward 5’-GCCCTCTCTCTCCTCTTGCT-3’,
>
> mCCL4 reverse 5’-CTGGTCTCATAGTAATCCATC-3’,
>
> mCCL5 forward 5’-GCCCACGTCAAGGAGTATTTCTA-3’
>
> and mCCL5 reverse 5’-ACACACTTGGCGGTTCCTTC-3’.

### Generation of VACV C7 specific polycolnal antibodies

The vaccinia C7L gene was cloned into bacterial expression vector-pET28-SUMO. The C7 expression plasmids were transfected into *E. coli* BL21 (DE3) cells. Cultures (2-liter) from a single transformant were grown at 37°C in LB Broth containing 100 µg/ml ampicillin until the *A*_600_ reached 0.6. The cultures were adjusted to 0.5 mM isopropyl-β-d-thiogalactopyranoside (IPTG), and then incubated for 20 h at 18°C with constant shaking. Cells were harvested by centrifugation and resuspended in buffer A (50 mM Tris–HCl, pH 7.5, 500 mM NaCl, 20 mM imidazole, 10% glycerol). The cells were lysed by sonication and the insoluble material was removed by centrifugation at 15,000 rpm for 45 min. The bacterial lysates were mixed for 1 h with 5 ml of Ni-NTA resin (Qiagen) that had been equilibrated with buffer A. The resins were poured into gravity-flow columns and then washed with 60 ml of buffer A. The adsorbed proteins were step-eluted with 300 mM imidazole in buffer A. The polypeptide compositions of the eluate fractions were monitored by SDS-PAGE and the peak fractions containing each recombinant protein were pooled. The eluates were dialyzed against buffer containing 50 mM Tris-HCl (pH 8), 200 mM NaCl, 2 mM DTT, 2 mM EDTA, 10% glycerol, and 0.1% Triton X-100 and then stored at –80 °C. Rabbit immunization ang generation of anti-C7 polyclonal rabbit antibody was performed in Pocono Rabbit Farm and Laboratory (PRF&L). The anti-C7 antibodies were purified from rabbit serum using affinity purification.

### Bone marrow chimeric mice experiments

Wild-type B6.SJL mice (CD45.1 background) or IFNAR1^−/−^ mice (CD45.2 background) were given a dose of irradiation (1096 Rads). After 6 hours, mice were injected retro-orbitally with isolated WT or KO bone marrow cells (5 × 10^6^ cells per mouse). Antibiotic-containing water (80 mg/L trimethoprim and 400 mg/L sulfamethoxazole) were provided for four weeks after irradiation. After another 4 weeks, the mice are ready for experiments.

### Statistics

Two-tailed unpaired Student’s t test was used for comparisons of two groups in the studies. Survival data were analyzed by log-rank (Mantel-Cox) test. The p values deemed significant are indicated in the figures as follows: *, p < 0.05; **, p < 0.01; ***, p < 0.001; ****, p < 0.0001. The numbers of animals included in the study are discussed in each figure legend.

