## Supplemental Figures for "Lung type II alveolar epithelial cells collaborate with CCR2^+^ inflammatory monocytes in host defense against an acute vaccinia infection in the lungs"

Ning Yang[1*], Joseph M. Luna[3], Peihong Dai[1,2], Yi Wang[1], Charles M. Rice[3], Stewart Shuman[2], and Liang Deng[1,3,4*]

### Figure S1

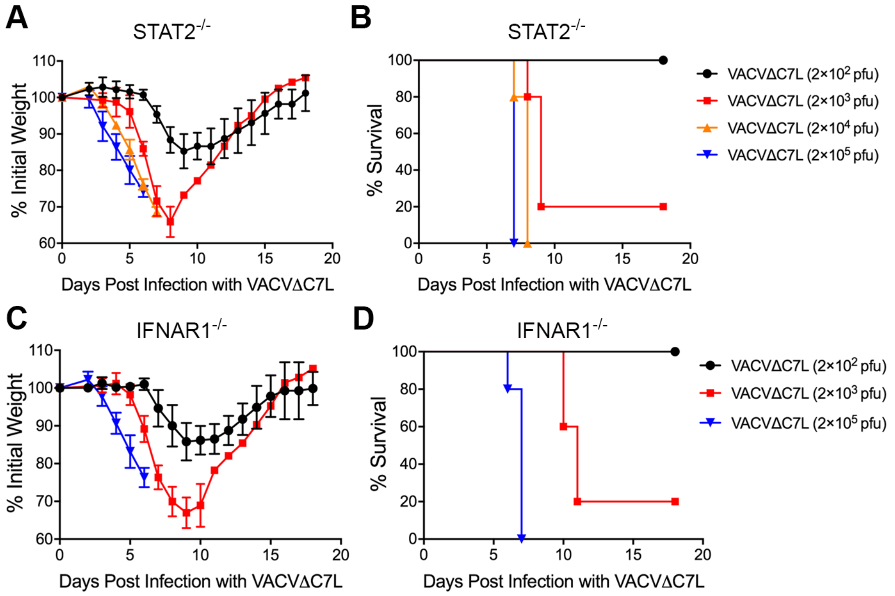
A
STAT2-/-
% Initial Weight
Days Post Infection with VACVΔC7L
B
STAT2-/-
% Survival
Days Post Infection with VACVΔC7L
VACVΔC7L (2×10^2 pfu)
VACVΔC7L (2×10^3 pfu)
VACVΔC7L (2×10^4 pfu)
VACVΔC7L (2×10^5 pfu)
C
IFNAR1-/-
% Initial Weight
Days Post Infection with VACVΔC7L
D
IFNAR1-/-
% Survival
Days Post Infection with VACVΔC7L
VACVΔC7L (2×10^2 pfu)
VACVΔC7L (2×10^3 pfu)
VACVΔC7L (2×10^5 pfu)

**Figure S1 related to Figure 1. Determination of LD50 of VACVΔC7L intranasal infection in STAT2$^{-/-}$ and IFNAR1$^{-/-}$ mice.** STAT2$^{-/-}$ mice were infected with VACVΔC7L at 2 x $10^2$ pfu, 2 x $10^3$ pfu, 2 x $10^4$ pfu, or 2 x $10^5$ pfu. IFNAR1$^{-/-}$ mice were infected with VACVΔC7L at 2 x $10^2$ pfu, 2 x $10^3$ pfu, or 2 x $10^5$ pfu. Mice were monitored for weight daily.
(A) and (C) shown are the percentages of initial weight over days in STAT1$^{-/-}$ mice (A) and IFNAR1$^{-/-}$ mice (C) post intranasal infection with VACVΔC7L at increasing doses.
(B) and (D) Kaplan-Meier survival curve of STAT2$^{-/-}$ (B) or IFNAR1$^{-/-}$ mice (D) infected with VACVΔC7L at increasing doses (n=5 in each group).

**Figure S2**

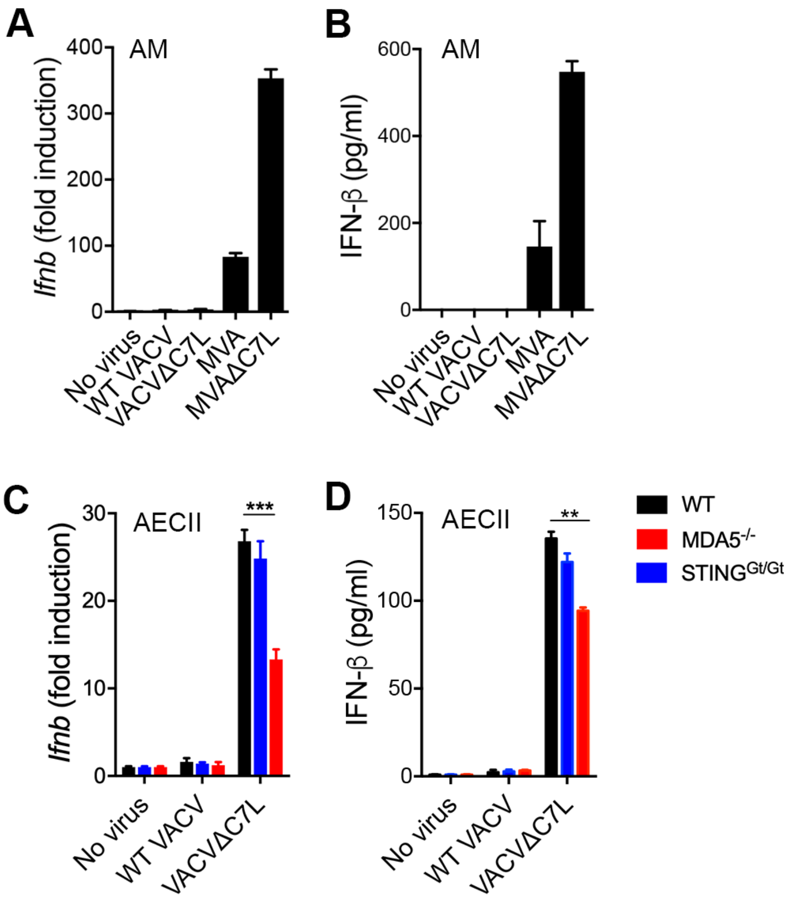
A
AM
Ifnb (fold induction)
400
300
200
100
0
No virus
WT VACV
VACVΔC7L
MVA
MVAΔC7L
B
AM
IFN-β (pg/ml)
600
400
200
0
No virus
WT VACV
VACVΔC7L
MVA
MVAΔC7L
C
AECII
Ifnb (fold induction)
30
20
10
0
***
No virus
WT VACV
VACVΔC7L
D
AECII
IFN-β (pg/ml)
150
100
50
0
**
No virus
WT VACV
VACVΔC7L
WT
MDA5-/-
STINGGt/Gt

**Figure S2 related to Figure 2. Infection of alveolar macrophages (AMs) in vitro with VACVΔC7L does not induce IFNB gene expression or IFN-β production.** AMs ($SiglecF^+CD11C^+$) were isolated from BAL of non-infected WT C57BL/6J mice. The cells were WT VACV, VACVΔC7L, MVA or MVAΔC7L at a MOI of 10. AECII were generated through culturing lineage negative epithelial progenitor cells isolated from WT, $STING^{Gt/Gt}$, or $MDA5^{-/-}$ mice as described in Figure 2. The cells were infected with either WT VACV or VACVΔC7L at a MOI of 10. The cells were collected at 12 h post infection and the supernatants were collected at 24 h post infection. RT-PCR analysis was performed to determine IFNB gene expression. ELISA analysis was performed to determine IFN-β levels in the supernatants of infected cells.

(A) RT-PCR analyses of Ifnb gene expression of AMs infected with either WT VACV, VACVΔC7L, MVA, or MVAΔC7L at a MOI of 10. Data are represented as mean ± SEM.

(B) ELISA analyses of IFN-β levels in the supernatants of AMs infected with either WT VACV, VACVΔC7L, MVA, or MVAΔC7L. Data are represented as mean ± SEM.

(C) RT-PCR analyses of Ifnb gene expression of AECIIs from WT, $STING^{Gt/Gt}$, or $MDA5^{-/-}$ mice infected with either WT VACV or VACVΔC7L. PBS was used as a mock infection control. Data are represented as mean ± SEM.

(B) ELISA analyses of IFN-β levels in the supernatants of AECIIs from WT, $STING^{Gt/Gt}$, or $MDA5^{-/-}$ mice infected with either WT VACV or VACVΔC7L. PBS was used as a mock infection control. Data are represented as mean ± SEM.

**Figure S3**

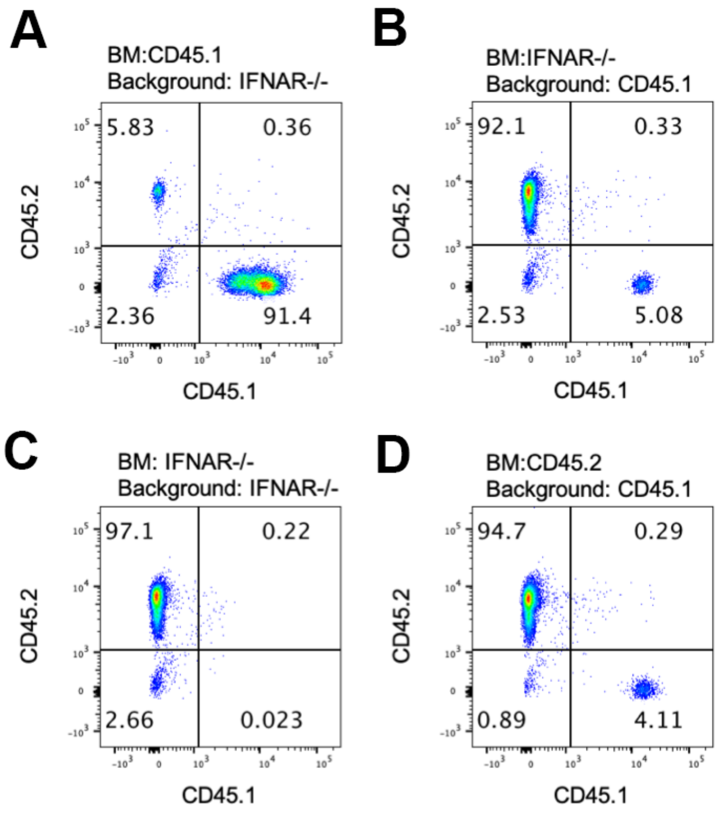
A
BM:CD45.1
Background: IFNAR-/-
5.83
0.36
2.36
91.4
CD45.2
CD45.1
B
BM:IFNAR-/-
Background: CD45.1
92.1
0.33
2.53
5.08
CD45.2
CD45.1
C
BM: IFNAR-/-
Background: IFNAR-/-
97.1
0.22
2.66
0.023
CD45.2
CD45.1
D
BM:CD45.2
Background: CD45.1
94.7
0.29
0.89
4.11
CD45.2
CD45.1

**Figure S3 is related to Figure 4. Analysis of CD45.1 and CD45.2 markers of immune cells in the bone marrow chimeras to confirm desired reconstitution of hematopoietic cells in the blood.**

(A) Dot plots of CD45.1 and CD45.2 staining of immune cells in the blood of reconstituted WT (CD45.1)→ IFNAR1$^{-/-}$ (CD45.2) mice, showing most of the CD45$^{+}$ cells carry CD45.1marker.

(B) Dot plots of CD45.1 and CD45.2 staining of immune cells in the blood of reconstituted IFNAR1$^{-/-}$ (CD45.2)→ WT (CD45.1) mice, showing most of the CD45$^{+}$ cells carry CD45.2 marker.

(C) Dot plots of CD45.1 and CD45.2 staining of immune cells in the blood of reconstituted IFNAR1$^{-/-}$ (CD45.2)→ IFNAR1$^{-/-}$ (CD45.2) mice, showing 97% of the CD45$^{+}$ cells carry CD45.2 marker.

(D) Dot plots of CD45.1 and CD45.2 staining of immune cells in the blood of reconstituted WT (CD45.2)→ WT (CD45.1) mice, showing most of the CD45$^{+}$ cells carry CD45.2 marker.

**Figure S4**

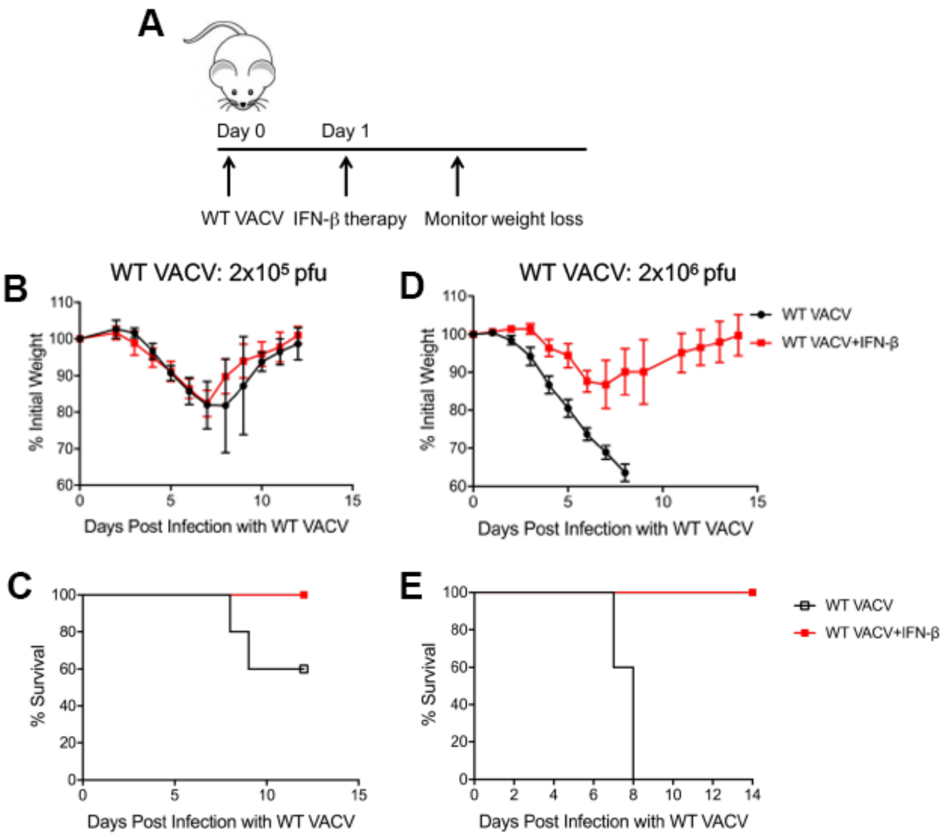
A
Day 0
Day 1
WT VACV
IFN-β therapy
Monitor weight loss
B
WT VACV: 2x10⁵ pfu
% Initial Weight
Days Post Infection with WT VACV
D
WT VACV: 2x10⁶ pfu
WT VACV
WT VACV+IFN-β
% Initial Weight
Days Post Infection with WT VACV
C
% Survival
Days Post Infection with WT VACV
E
% Survival
Days Post Infection with WT VACV
WT VACV
WT VACV+IFN-β

**Figure S4 related to Figure 4. Intranasal administration of IFN-β rescues mice from lethal WT VACV infection.**

(A) experimental scheme to test whether intranasal administration of IFN-β rescues mice from lethal infection of WT VACV. 6-8 week old WT C57BL/6J mice with WT VACV at 2 x $10^5$ pfu or 2 x $10^6$ pfu. They were either treated with intranasal administarion of IFN-β (1 μg per mouse) or PBS at day one post infection. Mice weight and survival were monitored daily.

(B) shown are the percentages of initial weight over days in mice infected with WT VACV at 2 x $10^5$ pfu followed by IFN-β treatment or PBS mock treatment.

(C) Kaplan-Meier survival curve of mice infected with WT VACV at 2 x $10^5$ pfu followed by IFN-β treatment or PBS mock treatment (n=5 in each group).

(D) shown are the percentages of initial weight over days in mice infected with WT VACV at 2 x $10^6$ pfu followed by IFN-β treatment or PBS mock treatment.

(E) Kaplan-Meier survival curve of mice infected with WT VACV at 2 x $10^6$ pfu followed by IFN-β treatment or PBS mock treatment (n=5 in each group).

**Figure S5**

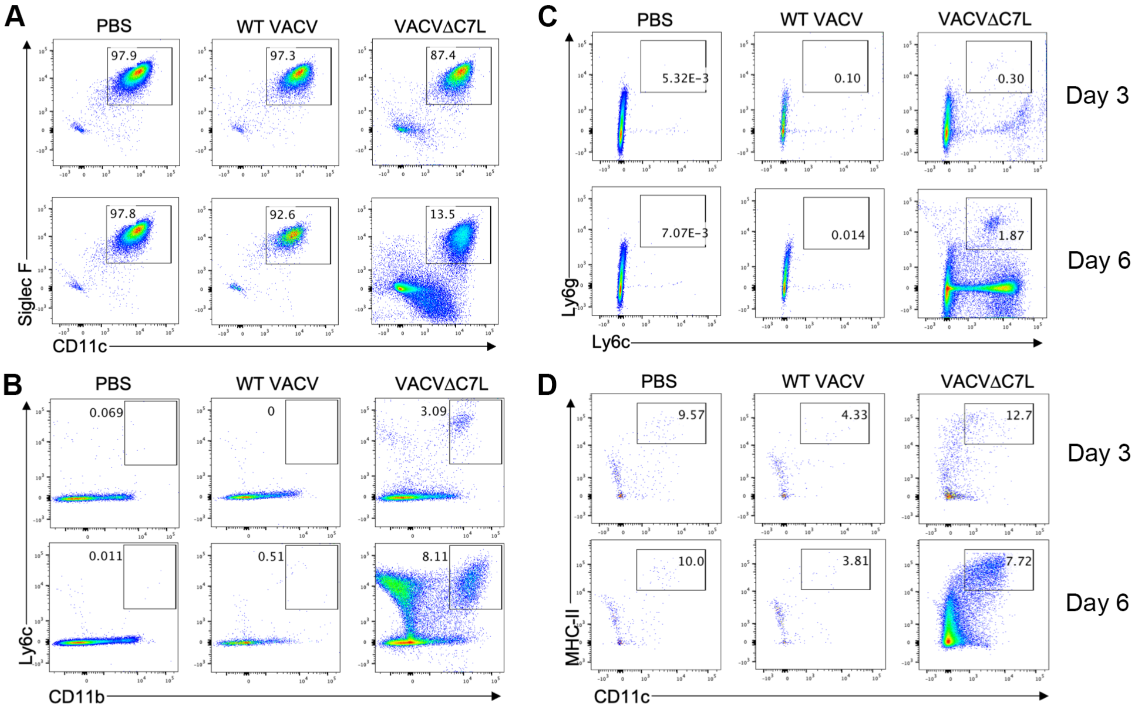
A
PBS
WT VACV
VACVΔC7L
97.9
97.3
87.4
97.8
92.6
13.5
Siglec F
CD11c
B
PBS
WT VACV
VACVΔC7L
0.069
0
3.09
0.011
0.51
8.11
Ly6c
CD11b
C
PBS
WT VACV
VACVΔC7L
5.32E-3
0.10
0.30
Day 3
7.07E-3
0.014
1.87
Day 6
Ly6g
Ly6c
D
PBS
WT VACV
VACVΔC7L
9.57
4.33
12.7
Day 3
10.0
3.81
7.72
Day 6
MHC-II
CD11c

**Figure S5 related to Figure 5. Intranasal infection of VACVΔC7L results in the influx of dendritic cells (DCs), monocytes, neutrophils, CD8$^+$, and CD4$^+$ T cells into bronchoalveolar space of the infected lungs.** WT C57BL/6J mice were infected with either WT VACV at 2 x $10^5$ pfu or with VACVΔC7L at 2 x $10^7$ pfu, or mock-infected with PBS. BAL was collected at 3 and 6 days post infection or PBS treatment. The myeloid cell populations in the BAL were analyzed by FACS.

(A) Dot plots of Siglec F$^+$CD11c$^+$ lung alveolar macrophages in the BAL from mice infected with either WT VACV, VACVΔC7L, or mock infected.

(B) Dot plots of Ly6C$^+$CD11b$^+$ inflammatory monocytes in the BAL from mice infected with either WT VACV, VACVΔC7L, or mock infected.

(C) Dot plots of Ly6G$^+$Ly6C$^+$ neutrophils in the BAL from mice infected with either WT VACV, VACVΔC7L, or mock infected.

(D) Dot plots of MHCII$^+$CD11c$^+$ DCs in the BAL from mice infected with either WT VACV, VACVΔC7L, or mock infected.
